# *Eomes* and *Brachyury* control pluripotency exit and germ layer segregation by changes of chromatin state

**DOI:** 10.1101/774232

**Authors:** Jelena Tosic, Gwang-Jin Kim, Mihael Pavlovic, Chiara M. Schröder, Sophie-Luise Mersiowsky, Margareta Barg, Alexis Hofherr, Simone Probst, Michael Köttgen, Lutz Hein, Sebastian J. Arnold

## Abstract

The first lineage specification of pluripotent mouse epiblast segregates neuroectoderm (NE) from mesoderm and endoderm (ME) by currently poorly understood mechanisms. Here we demonstrate that the induction of any ME-gene programs critically relies on the T-box (Tbx) transcription factors *Eomes* and *Brachyury* that concomitantly repress pluripotency and NE gene programs. Tbx-deficient cells retain pluripotency and differentiate to NE lineages despite the presence of ME-inducing signals TGFβ/Nodal and WNT. Pluripotency and NE gene networks are additionally repressed by Tbx-induced ME factors, demonstrating a remarkable redundancy in program regulation to safeguard mutually exclusive lineage specification. Chromatin analyses revealed that accessibility of ME-gene enhancers depends on Tbx-binding, while NE-gene enhancers are accessible and activation-primed already at pluripotency state. This asymmetry of chromatin landscape thus explains the default differentiation of pluripotent cells to NE in the absence of ME-induction mediated through the activating and repressive functions of early Tbx factors *Eome*s and *Brachyury*.

---

The first cell lineage specification of the embryo proper that segregates neuroectoderm (NE) from mesoderm and definitive endoderm (ME) takes place in the epiblast (EPI) of the postimplantation mouse embryo^1^. EPI cells reside in a primed pluripotent state^2, 3^ that follows ground state pluripotency of the inner cell mass established in the embryonic day 3.5 (E3.5) blastocyst^4^. Around E6.5 rising levels of extracellular signaling molecules TGFβ and WNT on the prospective posterior side of the embryo^1^ pattern the EPI and instruct transcriptional identities, thus promoting pluripotency exit and specification of the three germ layers: definitive endoderm, mesoderm, and NE^5^.

Different subpopulations of ME lineages are induced by TGFβ/Nodal and WNT signals and their respective downstream target genes including T-box (Tbx) transcription factors, *Eomesodermin* (*Eomes*) and *Brachyury* (*T*)^1^. Anterior mesoderm and definitive endoderm specification critically depends on functions of *Nodal*-induced *Eomes*, which is expressed in the posterior EPI between E6.5 until E7.5^6–9^. *Wnt3/Wnt3a*-induced *Brachyury* expression partially overlaps with *Eomes* expression in the posterior EPI and primitive streak, however, *Brachyury*-deficient embryos show little defects in the anterior mesoderm, but fail to extend posteriorly and lack axial midline mesoderm^10, 11^. Despite the phenotypic differences between embryos lacking either *Eomes* or *Brachyury*, molecular studies indicate largely overlapping targets when assessing genome-wide chromatin binding and transcriptional responses^8, 12–17^. Thus, important aspects of the unique and redundant functions of *Eomes* and *Brachyury* have yet to be characterized. The prevailing model of NE induction suggests that neural specification occurs as a default path of differentiation, in the absence of ME-inducing signals^18–21^. NE formation in the mouse takes place in the anterior epiblast, where ME-inducing WNT3/WNT3A and NODAL signals are antagonized by LEFTY1, CER1, and DKK1 secreted by the anterior visceral endoderm (AVE)^22–25^. Next, intrinsic regulatory networks of transcription factors such as *Sox2*, *Pou3f1* (alias *Oct6*), *Zeb2*, *Otx2, Zic2, and Zic3* reinforce neural fate by regulating the expression of NE gene programs and repressing pluripotency and ME genes^26–28^. Despite the conceptual framework of ME induction by extracellular signals, and NE differentiation as default in the absence of ME-inducing signals, the precise transcriptional and epigenetic mechanisms of this mutually exclusive lineage choice remain unexplored.

In this study we reveal essential roles of the Tbx factors *Eomes* and *Brachyury* for the exit from pluripotency during the segregation of the NE and ME lineages downstream of TGFβ/WNT signals. The activation of any ME gene program critically depends on transcriptional control by either of these two Tbx factors. Cells deficient for both Tbx factors (dKO) fail to differentiate to ME and retain primed pluripotency in ME-inducing conditions. After prolonged culture dKO cells differentiate to neurons despite the presence of ME-inducing/NE-blocking signals, TGFβ and WNT. We demonstrate that *Eomes* and *Brachyury* directly bind and repress enhancers of pluripotency and NE-associated genes that possess open chromatin conformation in pluripotent cells. In contrast, chromatin accessibility of ME enhancers depends on Tbx factor activities and ME enhancers remain in an inaccessible state in Tbx-deficient cells. Additionally, cascades of Tbx-induced ME-specifying transcription factors contribute to the repression of pluripotency and NE gene programs. In conclusion, we elucidate the mechanisms of lineage segregation of NE and ME cell types involving changes in chromatin landscape and transcriptional activation and repression, which are directly mediated by Tbx factors and indirectly by Tbx-induced ME transcription factors.

## Results

### *Eomes* and *Brachyury* specify all mesoderm and endoderm lineages

To study transcriptional regulation of pluripotency exit and early lineage determination we genetically deleted Tbx factors *Brachyury* and *Eomes* in mouse embryonic stem cells (ESCs) and investigated their differentiation potential. Briefly, we generated ESCs carrying fluorescent reporters in the start codon of one allele (*Eomes*^Gfp^ and *Bra*^Tom^) and frame-shift deletions in exon 1 of the other allele to homozygously delete *Brachyury* (BraKO) and *Eomes* (EoKO) individually or as double knock-out (dKO) cells (Fig. 1a; Supplementary Fig. 1a). ESCs differentiation towards ME lineages was induced by embryoid body (EB) formation in serum-free medium and administration of the NODAL analog ActivinA (ActA, Fig. 1b)^29^. Time-course expression analysis in WT cells indicates that *Eomes* and *Brachyury* expression shortly precedes other ME-marker genes (Supplementary Fig. 1b). The absence of *Brachyury* and *Eomes* mRNA and protein was confirmed by qRT-PCR and immunofluorescence (IF) staining in individual KO and dKO cells (Supplementary Fig. 1c, d). Abundant Gfp and Tom reporter activation was observed from day 3 of differentiation in dKO cells (Supplementary Fig. 1e), indicating the competence of WT and dKO cells to respond to WNT and TGFβ signals. This signaling competence was confirmed by Western blot analysis for non-phosphorylated (active) β-CATENIN and phosphorylated SMAD2^30–33^ (Supplementary Fig. 1f,g), and by reporter assays using WNT- and TGFβ- responsive luciferase reporters (Supplementary Fig. 1h,i).

**Fig. 1:**
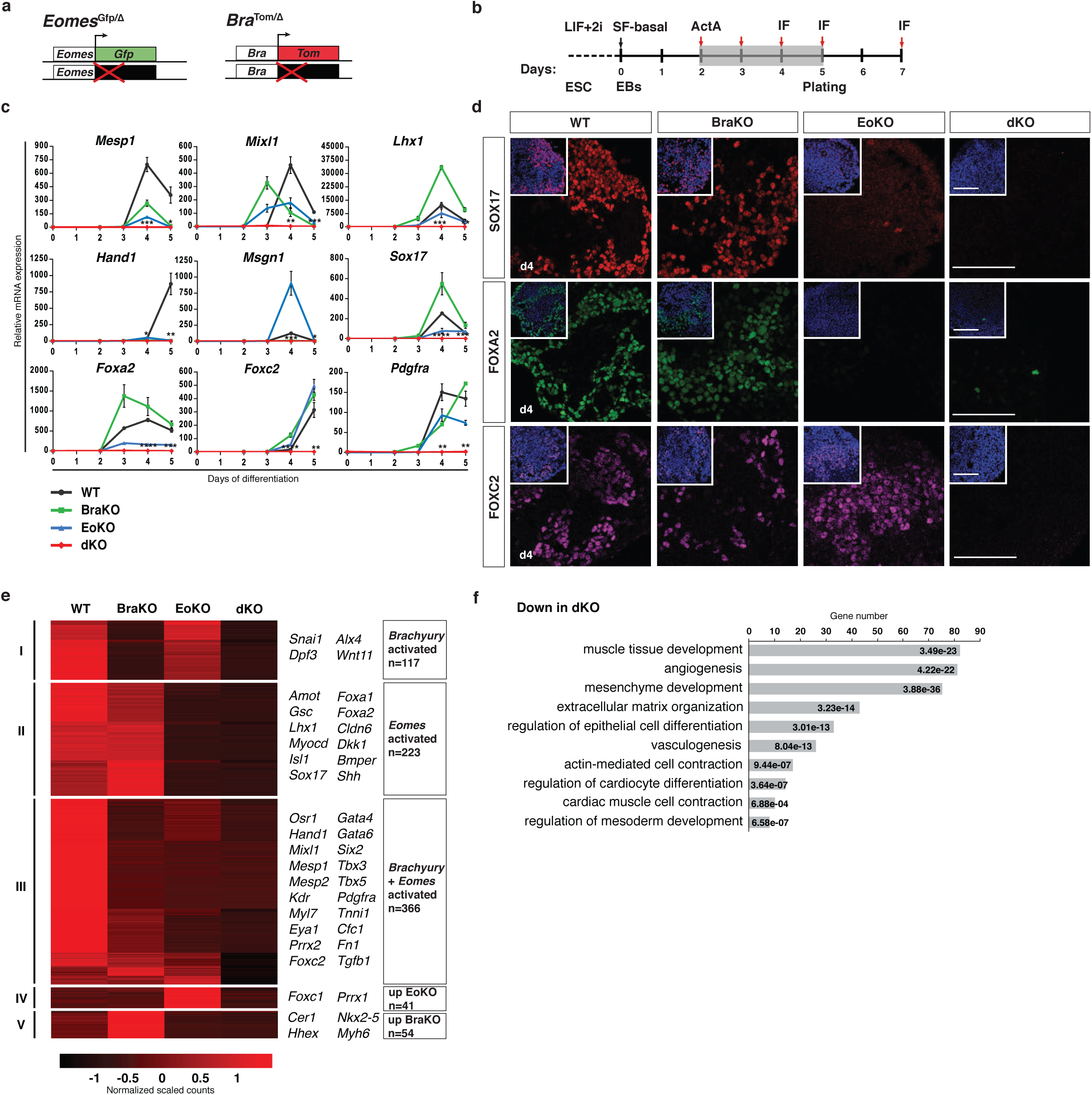
*Eomes-* and *Brachyury*-deficient cells fail to form any type of mesoderm and definitive endoderm. **a**, Schematic of *Eomes*^Gfp/Δ^ (EoKO) and *Bra*^Tom/Δ^ (BraKO) loss-of-function alleles in mouse embryonic stem cells (ESCs). One allele contains a fluorescent reporter (*Gfp* or *Tomato*, *Tom*) knock-in, the second allele contains frame-shift deletions in exon 1 of each gene. **b**, Differentiation protocol to mesoderm and definitive endoderm (ME) cell types: mouse ESCs are maintained in LIF+2i medium, followed by aggregation to embryoid bodies (EBs) in serum-free (SF-basal) medium for 2 days, and treatment with ActivinA (ActA) from day 2 to day 5. Arrows indicate time points of analysis. **c**, Relative mRNA expression of ME markers during differentiation of WT, BraKO, EoKO, and double knock-out (dKO) cells measured by qRT-PCR in a 5-day time-course showing the complete absence of ME differentiation in dKO cells. Error bars indicate SEM. p-values for differences of mean expression between WT and dKO samples were calculated by Student’s t-test. *:p≤0.05; **:p≤0.01; ***: p≤0.001; ****: p≤0.0001. **d**, Immunofluorescence (IF) staining of definitive endoderm (SOX17 and FOXA2) and mesoderm (FOXC2) markers at day 4 of differentiation. Scale bars 100 μm. **e**, Heatmap showing clustered groups (I to V) of downregulated genes (adjusted p-value≤0.05, log_2_(FC)≤-2.5) in dKO compared to WT cells at day 5 of differentiation analyzed by RNA-seq. Scale indicates centered scaled counts normalized by library size and gene-wise dispersion. Important genes in each group are indicated to the right. **f**, Gene ontology (GO) enrichment analysis of downregulated genes in dKO compared to WT cells indicating the significantly reduced signatures of ME developmental programs in dKO cells. p-values for each term are indicated.

To evaluate the differentiation potential of BraKO, EoKO, and dKO cells, we monitored mRNA expression of markers for anterior mesoderm (*Mesp1*), early mesendoderm (*Mixl1, Lhx1*), paraxial and lateral plate mesoderm (*Hand1, Msgn1*), definitive endoderm (*Sox17*, *Foxa2*) and pan-mesoderm (*Foxc2, Pdgfra*) during a 5-day time-course of differentiation by qRT-PCR (Fig. 1c). As expected, BraKO cells fail to express the target gene *Msgn1*^34^, EoKO cells show reduced *Lhx1, Sox17,* and *Foxa2*^8, 17, 35^ levels, while the expression of *Brachyury* and *Eomes* co-regulated target genes *Mesp1, Mixl1,* and *Hand1* is decreased in both BraKO and EoKO compared to WT cells. Remarkably, dKO cells fail to express any of the tested ME markers, including *Foxc2* and *Pdgfra* transcripts that remain expressed in individual KO cells, thus indicating that ME differentiation is abolished in dKO cells (Fig. 1c). This observation was confirmed by IF staining for endoderm markers SOX17 and FOXA2 and mesoderm markers FOXC2 and FIBRONECTIN1 (FN1, Fig. 1d; Supplementary Fig. 1j). To globally analyze the differentiation potential of dKO cells we performed transcriptional profiling by RNA sequencing (RNA-seq) at day 5 of differentiation. 801 significantly downregulated genes in dKO compared to WT cells were clustered in 5 distinct groups (Fig. 1e; Supplementary Table 1). Clusters correspond to *Brachyury*-activated (cluster I), *Eomes*-activated (cluster II), and *Brachyury* and *Eomes* co-regulated genes (cluster III) that are reduced or absent in dKO cells. Clusters IV and V contain inversely regulated genes by *Brachyury* and *Eomes*, such that their expression is increased above WT levels in EoKO, or in BraKO respectively (Fig. 1e). Importantly, differentially downregulated genes confirmed the complete absence of the ME gene signature in dKO cells, underscored by gene ontology (GO) analysis revealing gene functions in ME-derived organ formation such as cardiac, angiogenic, and muscle development (Fig. 1f; Supplementary Table 1). In summary, the absence of both Tbx factors *Brachyury* and *Eomes* abrogates formation of any ME lineages despite responsiveness to TGFβ and WNT signals.

### Tbx-deficient cells retain primed pluripotency and differentiate into neurons

Next, we analyzed the transcriptional signature of the 429 upregulated genes in dKO compared to WT cells that were clustered in 3 distinct groups (Fig. 2a; Supplementary Table 2). Cluster I contains transcripts upregulated in BraKO and dKO cells, including several neural markers (*Nkx6-2*, *Neurog2,* and *Pax6*). Cluster II represents genes upregulated in EoKO and dKO cells, such as *Epha2*, *Zic1,* and *Hoxa2* marking the anterior EPI and NE. The largest number of genes show significantly increased expression only in dKO cells (cluster III, n=212) and represent pluripotency- and epiblast stem cell (EpiSC)-associated genes *Pou5f1* (alias *Oct4*), *Nanog*, *Sox2*, *Esrrb, Wnt8a, Lefty2, Nkx1-2*, and early neural markers, such as *Sox1, Sox3, Pou3f1, Olig3,* and *Neurod1* (Fig. 2a). GO-term analysis shows enrichment of functions associated with neurons and stem cells in differentiated dKO cells (Fig. 2b; Supplementary Table 2). We compared all upregulated transcripts in dKO cells with expression signatures of NE cells generated from ESCs by BMP and TGFβ signaling inhibition^36^ and of EpiSCs^37^ and found remarkable overlap with NE (n=251) and EpiSCs (n=97) marker genes (Fig. 2c; Supplementary Table 2). qRT-PCR expression analysis of core pluripotency (*Pou5f1, Nanog,* and *Sox2*), and EpiSC markers (*Wnt8a, Lefty2,* and *Nkx1-2)* during a 5-day time-course of differentiation confirmed the maintenance of primed pluripotency in dKO cells, while they properly exit from ground state pluripotency shown by the downregulation of *Zfp42* (alias *Rex1;* Fig. 2d)^2, 38^. Following the expression dynamics of pluripotency genes alongside with *Eomes* and *Brachyury* transcripts revealed that the downregulation of pluripotency genes indeed coincides with the onset of expression of Tbx factors in WT cells (Supplementary Fig. 2a). qRT-PCR results showing increased pluripotency gene expression were confirmed by IF staining for pluripotency factors NANOG, SOX2, and OCT4 that show abundant nuclear staining in dKO cells at day 5 (Fig. 2e) and day 7 (Supplementary Fig. 2b) of differentiation, but are absent in WT, BraKO, and EoKO cells.

**Fig. 2:**
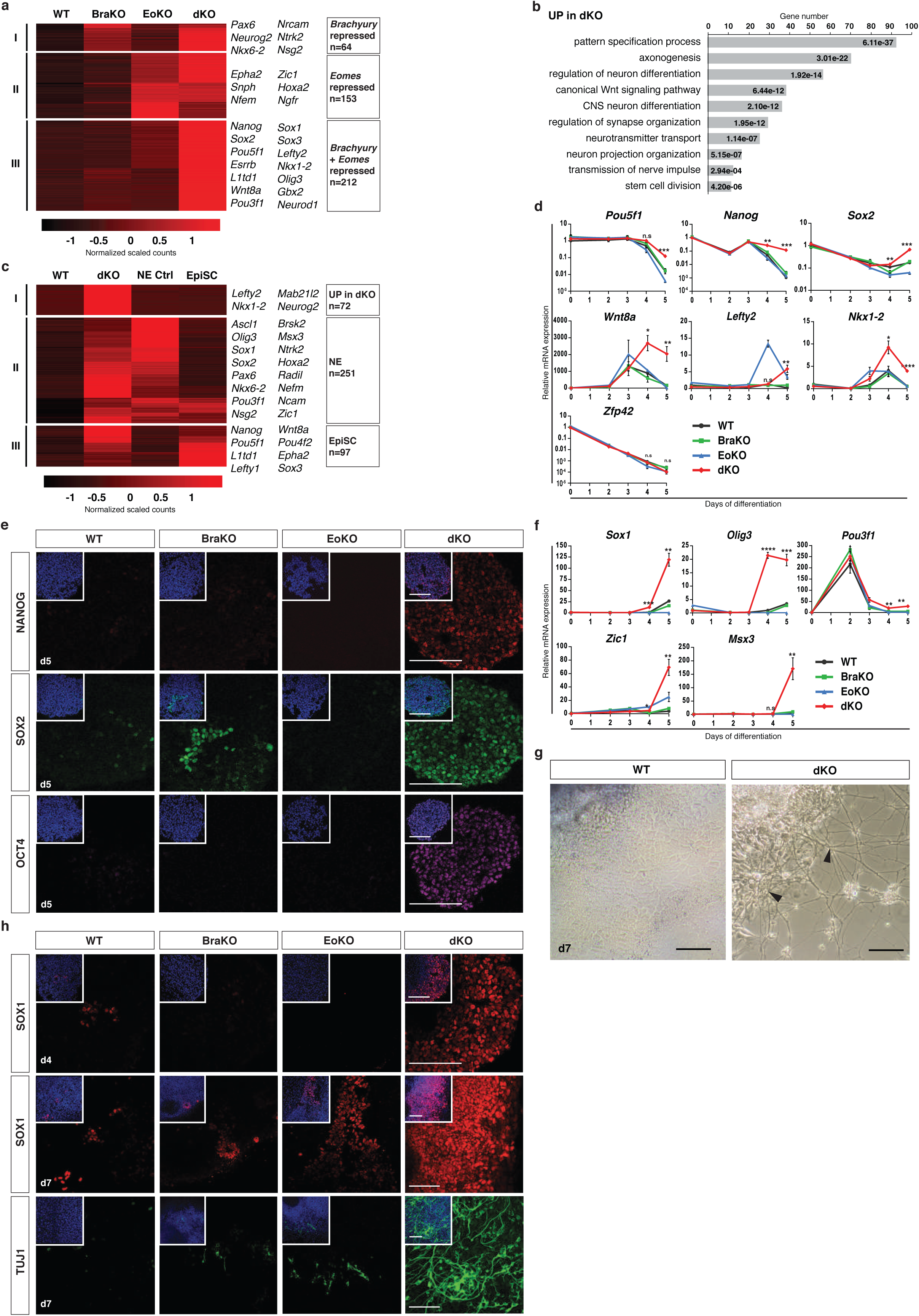
Tbx-deficient cells retain primed pluripotency and differentiate to neuroectoderm in ME-inducing conditions. **a**, Heatmap showing clustered groups (I to III) of upregulated genes (adjusted p-value≤0.05, log_2_(FC)≥2.5) in dKO compared to WT cells at day 5 of differentiation analyzed by RNA-seq. Representative genes within groups are indicated to the right. Scale indicates centered scaled counts normalized by library size and gene-wise dispersion in a and c. **b**, GO-term enrichment analysis of upregulated genes in a, indicating pluripotency and neuroectoderm (NE) expression signatures in dKO cells. The p-values for each term are indicated. **c**, Heatmap comparing the expression of upregulated genes in dKO (adjusted p-value≤0.05, log_2_(FC)≥2.5) with RNA-seq profiles of ESCs differentiated to NE (NE Ctrl) and epiblast stem cells (EpiSCs)^37^. Important epiblast (EPI) and NE markers are indicated to the right. **d**, Relative mRNA expression of markers for naïve (*Pou5f1*, *Nanog, Sox2,* and *Zfp42*) and primed pluripotency/EpiSCs (*Wnt8a, Lefty2,* and *Nkx1-2)* during 5 days of differentiation measured by qRT-PCR. Error bars indicate SEM; p-values for differences of mean expression between WT and dKO samples were calculated by Student’s t-test. n.s:p>0.05; *:p≤0.05; **:p≤0.01; ***: p≤0.001; ****: p≤0.0001 in d and f. **e**, IF staining for pluripotency markers NANOG, SOX2, and OCT4 at day 5 of differentiation indicating maintenance of pluripotency in dKO, but not in other cells. Scale bars 100 μm. **f**, Relative mRNA expression of anterior EPI and NE markers (*Sox1, Olig3, Pou3f1, Zic1,* and *Msx3*) during 5 days of differentiation measured by qRT-PCR. **g**, Bright field images of plated cells at day 7 of differentiation showing neuronal cell morphology with axonal processes (arrowheads) present in dKO, but not in WT cells. Scale bars 200 μm. **h**, IF staining for SOX1 at day 4, and SOX1 and TUJ1 at day 7 of differentiation indicating prominent neural marker staining in dKO, that is only occasionally observed in BraKO, EoKO, and WT cells. Scale bars 100 μm.

In addition, we monitored expression of early (*Sox1, Olig3,* and *Pou3f1*) and mature (*Zic1* and *Msx3*) NE markers during differentiation by qRT-PCR. These genes are significantly upregulated in dKO cells after 5 days of differentiation, but are absent in BraKO, EoKO, and WT cells (Fig. 2f). Expression of *Pou3f1* is transiently upregulated at day 2 of differentiation in WT cells and is downregulated during initiation of *Eomes* and *Brachyury* expression, while other NE markers are not expressed in WT cells under these experimental conditions (Supplementary Fig. 2c). Remarkably, at day 7 the differentiated dKO cells exhibit neuronal morphology, such as characteristic neuronal processes that are not found in WT cells (Fig. 2g). Immunostaining against neural markers SOX1 and TUJ1 (β-III-tubulin) confirmed neural differentiation of dKO cells, while only very few SOX1 and TUJ1 positive cells are detected in BraKO, EoKO, or WT cells (Fig. 2h). We performed double IF staining for OCT4/SOX1 and NANOG/SOX1 to reveal if NE and pluripotency markers are simultaneously expressed in dKO cells or arise from heterogenous cell population (Supplementary Fig. 2d,e). While the majority of dKO cells express only the neural marker SOX1, patches of OCT4+/SOX1- and OCT4+/SOX1+ cells are also detected (Supplementary Fig. 2d). Similarly, SOX1/NANOG staining resulted in a heterogeneous pattern with single and double positive cells (Supplementary Fig. 2e).

In conclusion, *Eomes* and *Brachyury* are not only crucial regulators for ME specification, but are also required to exit primed pluripotency and repress the default NE gene program during ME formation downstream of TGFβ/WNT signals.

### *T*^2J/2J^;*Eomes*^ΔEpi^ embryos show increased pluripotency and anterior neuroectoderm gene expression

We asked if observed phenotypes of differentiating dKO cells reflect embryonic functions of *Eomes* and *Brachyury* during gastrulation, when partial co-expression of both factors occurs in the posterior epiblast and primitive streak (Supplementary Fig. 3a). We analyzed embryos with epiblast-specific deletion of *Eomes* (*Eomes*^ΔEpi^)^7^ and homozygous deletion of *Brachyury* (*T*^2J/2J^)^10^ individually and as double mutants (*T*^2J/2J^;*Eomes*^ΔEpi^, hereafter dKO embryos). RNA-seq analysis of E7.5 epiblasts of *T*^2J/2J^ embryos shows little differences in the expression profiles compared to WT (Fig. 3a; Supplementary Table 3). Among the few genes that are reduced in *T*^2J/2J^ and dKO embryos are described BRACHYURY targets *Msgn1* and *Snai1* (cluster I)^13, 39^. Concordant with the more severe phenotype of *Eomes*^ΔEpi^ embryos at E7.5^7^, they show more differences in gene expression, including absence of early cardiac mesoderm (*Myl7, Myocd*, and *Tnnt2*)^8^ and definitive endoderm markers (*Cer1*, *Foxa2*, and *Gata6*)^7, 17^ (cluster II, Fig. 3a). In contrast, cluster III contains a large group of mesoderm markers that are abundantly expressed in either *T*^2J/2J^ or *Eomes*^ΔEpi^ embryos, but absent in dKO embryos, including *Pdgfra, Osr1, Tbx6,* and *Hand1* (Fig. 3a). The failure to specify ME is further confirmed by the GO-term analysis of downregulated genes in dKO embryos (Fig. 3b; Supplementary Table 3). Cluster analysis of upregulated genes in dKO embryos revealed a large group of EpiSC markers (*Lefty1/2*, *Tdgf1*, and *Evx1*) and anterior EPI and early NE genes (*Utf1*, *Epha2*, *Grik3*, *Zic5*, and *Slc7a3*) upregulated both in *Eomes*^ΔEpi^ and dKO embryos (cluster V, n=124, Fig. 3a). Similar to differentiating dKO cells, genes with strongly enhanced expression specifically in dKO embryos include pluripotency genes *Pou5f1*, *Nanog,* and *Esrrb*, and NE markers *Olig2, Zic2,* and *Mab21l2* (cluster VI, n=90, Fig. 3a). GO-term analysis of upregulated genes in dKO embryos confirmed the signature corresponding to neural differentiation (Fig. 3c; Supplementary Table 3), however, less prominent compared to dKO cells, probably due to variances in differentiation kinetics and the analyzed timepoints.

**Fig. 3:**
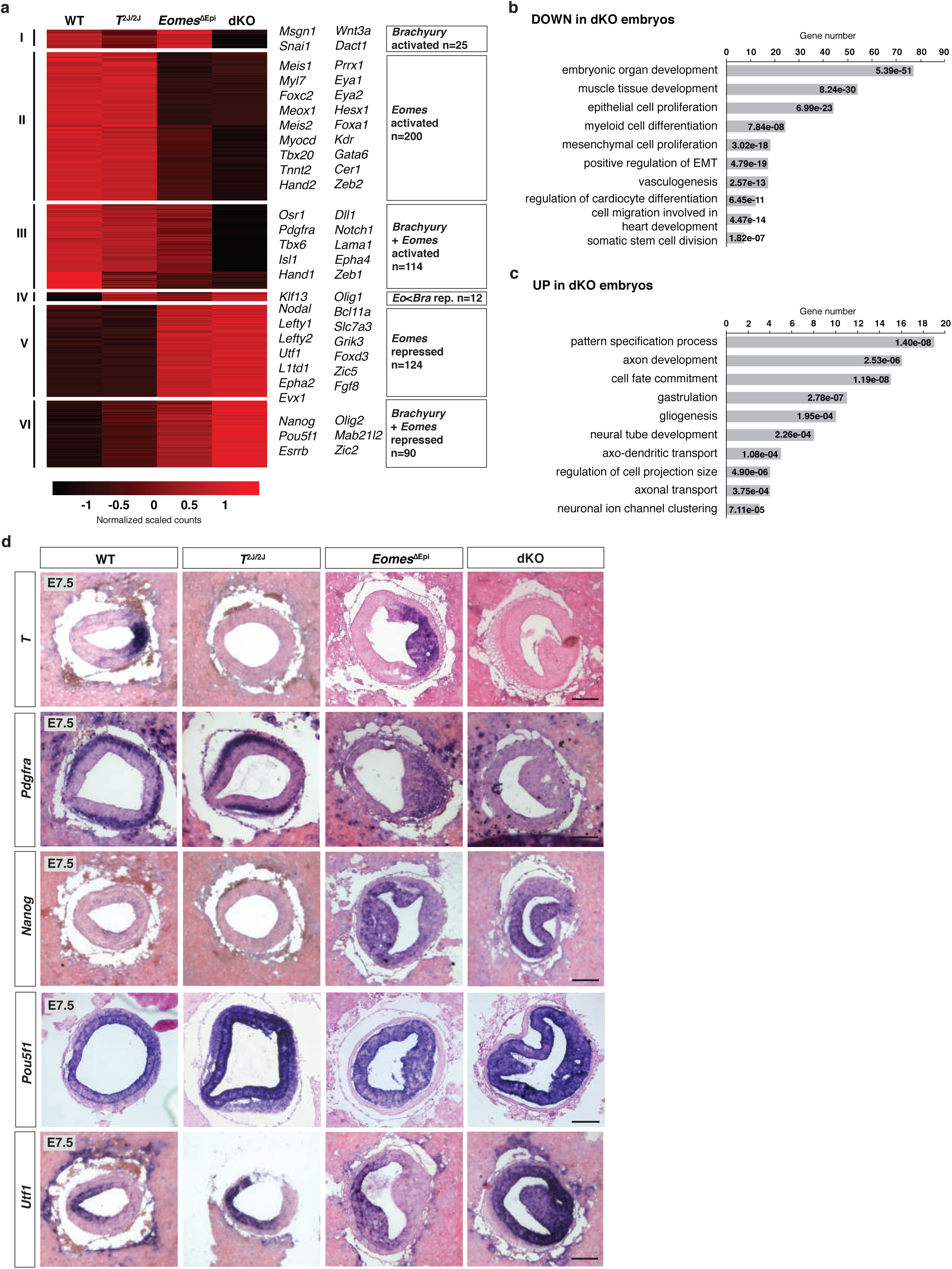
*Eomes* and *Brachyury* dKO embryos fail to form ME, maintain pluripotency and show increased expression of anterior EPI markers. **a**, Heatmap showing clusters (I to VI) of differentially expressed genes (adjusted p-value≤0.05, log_2_(FC) of +/-1.0) of *T*^2J/2J^, *Eomes*^ΔEpi^, and dKO (*T*^2J/2J^;*Eomes*^ΔEpi^) embryos compared to WT assayed by RNA-seq at E7.5. Representative markers are indicated to the right. Scale represents centered scaled counts normalized by library size and gene-wise dispersion. **b**, GO-term enrichment analysis of significantly downregulated genes in dKO compared to WT embryos showing overrepresentation of terms associated with ME development. The p-values for each term are shown in b and c. **c**, GO-term enrichment analysis of genes significantly upregulated in dKO embryos indicating processes related to neural development. **d**, mRNA *in situ* hybridization analysis of transversal sections of E7.5 embryos for *T* (*Brachyury*)*, Pdgfra, Nanog, Pou5f1,* and *Utf1* showing absence of mesoderm (*Pdgfra*) and expanded expression of pluripotency (*Nanog* and *Pou5f1*) and anterior epiblast (*Utf1*) markers in the dKO epiblast. Scale bars 100 μm.

To evaluate the expression patterns of some differentially regulated genes in dKO embryos we performed *in situ* hybridization on sections of E7.5 WT, *T*^2J/2J^, *Eomes*^ΔEpi^, and dKO embryos (Fig. 3d). The *T*^2J/2J^ homozygous genotype was inferred by the absence of *T* (*Brachyury)* mRNA, and *Eomes*^ΔEpi^ by the characteristic cell accumulations in the EPI^7^. *Pdgfra* is widely expressed in the mesoderm of WT, *T*^2J/2J^, and in the EPI of *Eomes*^ΔEpi^ embryos, but absent in dKO embryos (Fig. 3d). In contrast, pluripotency markers *Nanog* and *Pou5f1* and the anterior EPI/NE marker *Utf1* show expanded and/or enhanced expression in the entire EPI of dKO embryos. Interestingly, *T* and *Utf1* expression seem mutually exclusive in *Eomes*^ΔEpi^ embryos where the enlarged *T* expression domain restricts *Utf1* transcripts to the anterior (Fig. 3d). The expression of *Brachyury* target gene *Snai1*^13^ is completely lacking in *T*^2J/2J^ and dKO embryos, but it is increased in the *Eomes*^ΔEpi^ embryos (Supplementary Fig. 3b). Expression of the DE marker *Sox17* cannot be detected in *Eomes*^ΔEpi^ and dKO embryos, reflecting the DE specification defect, but is expressed within the endoderm layer of WT and *T*^2J/2J^ embryos (Supplementary Fig. 3b). In conclusion, dKO embryos and cells show similar phenotypes, namely the absence of all ME lineages and increased expression of pluripotency, anterior EPI and NE markers.

### Tbx factors bind and remodel enhancers of induced ME and repressed pluripotency and NE genes

To investigate the transcriptional regulation by EOMES and BRACHYURY on a genome-wide scale we analyzed chromatin occupancy and accessibility in WT and dKO cells. We performed chromatin immunoprecipitation and sequencing (ChIP-seq) of 5 day differentiated dKO cells with doxycycline (dox)-inducible GFP-tagged EOMES and BRACHYURY (EoGFP and BraGFP) that were extensively validated to recapitulate endogenous protein function (Supplementary Fig. 4a-j). Accordingly, the induced expression of EoGFP or BraGFP rescues the phenotype of dKO cells to very similar levels as the expression of full-length cDNAs of *Eomes* or *Brachyury* indicated by the close proximity to BraKO and EoKO cells, respectively, using principal component analysis (PCA) of RNA-seq data (Supplementary Fig. 4e). Using dKO+BraGFP and dKO+EoGFP cells for ChIP-seq identified 27,599 and 30,145 regions occupied by BRACHYURY and EOMES, respectively, which overlapped on 12,517 region-associated genes (Supplementary Fig. 4k; Supplementary Table 4). Due to the large overlap in target gene occupancy, we merged EOMES and BRACHYURY ChIP-seq data for the analyses of shared Tbx functions causing the dKO phenotype. Tbx-binding is frequently found in promoters (≤1 kb to gene bodies, 22%), introns (32%) and distal intergenic regions (34%), as expected for the binding of transcription factors (Supplementary Fig. 4l). Intersecting Tbx-bound regions (ChIP-seq) with differentially downregulated genes in dKO cells revealed 517 Tbx-occupied genes, including known ME-specifying factors (Fig. 4a; Supplementary Table 4). Importantly, the regulatory elements of 323 upregulated genes in dKO cells were also occupied by EOMES and BRACHYURY, including pluripotency (e.g. *Pou5f1, Nanog, Sox2, Esrrb*) and NE genes (e.g. *Pou3f1, Sox1, Zic1, Nkx6-2*) (Fig. 4b; Supplementary Table 4), suggesting functions of EOMES and BRACHYURY as transcriptional repressors of pluripotency and NE programs.

**Fig. 4:**
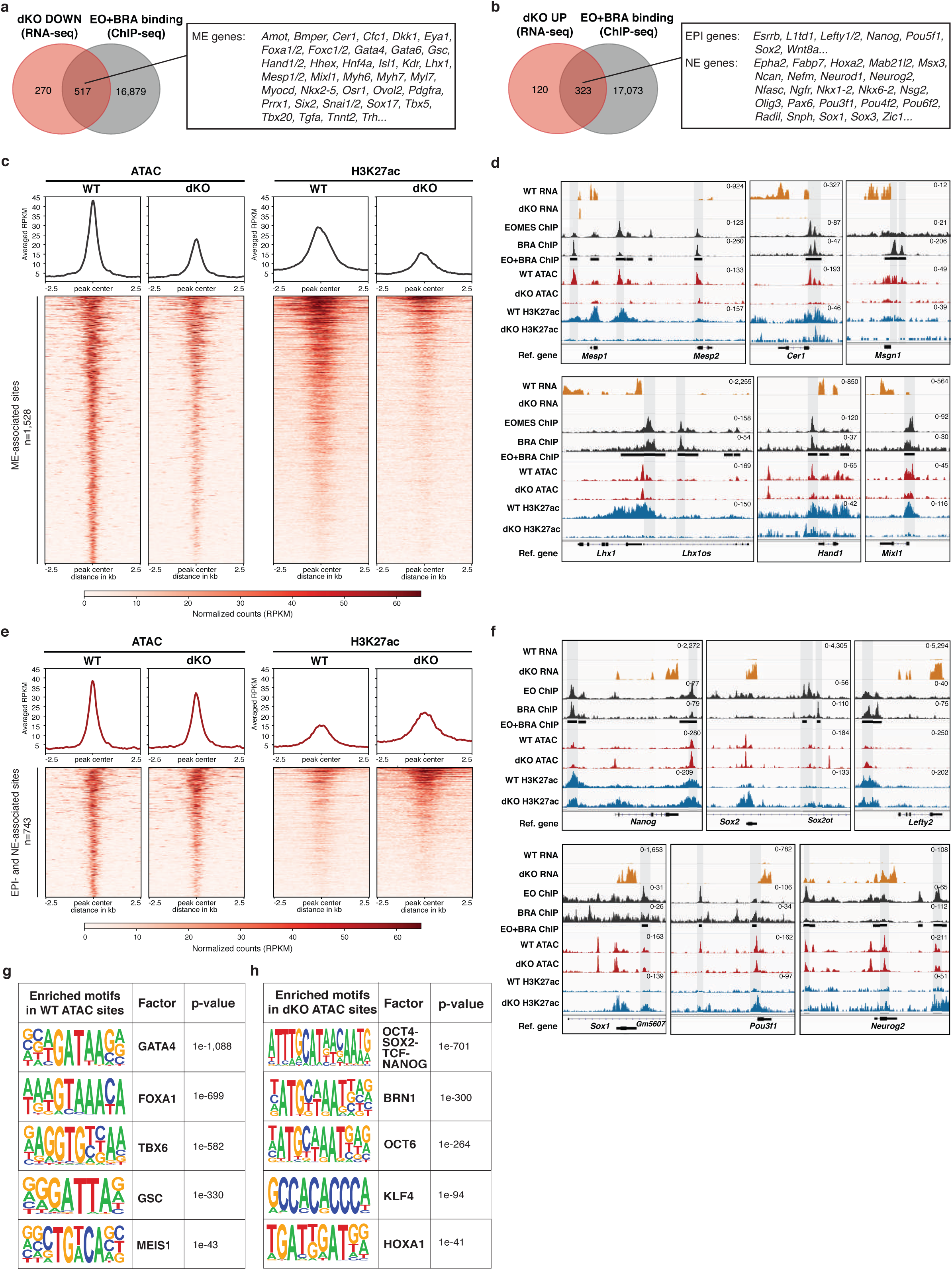
Tbx factors occupy enhancers of activated and repressed targets and remodel chromatin accessibility. **a**, Overlap of downregulated genes in differentiated dKO cells (RNA-seq) and genes associated with EOMES- or BRACHYURY-occupied regions (ChIP-seq). Out of 517 overlapping genes important ME regulators are indicated. **b**, Overlap of upregulated genes in differentiated dKO cells (RNA-seq) and genes associated with EOMES- or BRACHYURY-occupied regions (ChIP-seq). Out of 323 Tbx-repressed targets, markers of the EPI and NE are indicated. **c**, Heatmap of chromatin accessibility (ATAC-seq) and corresponding H3K27ac ChIP-seq signals of WT and dKO cells at EOMES- and BRACHYURY- occupied regions that are associated to downregulated genes in dKO showing reduced chromatin accessibility and H3K27ac marks in dKO cells. Scale represents normalized counts (reads per kilobase per million, RPKM) for H3K27ac ChIP-seq and ATAC-seq peak signals +/- 2.5 kb around the center of the peak in c and e. **d**, RNA-seq, ChIP-seq, and ATAC-seq coverage tracks for EOMES- and BRACHYURY-activated target genes *Mesp1, Cer1, Msgn1, Lhx1, Hand1, and Mixl1*. Counts normalized to RPKM are indicated in d and f. **e**, Heatmap of chromatin accessibility at Tbx-bound regions that are associated to upregulated genes in dKO, and corresponding H3K27ac ChIP-seq signals in WT and dKO cells. Chromatin accessibility is comparable between WT and dKO cells, while H3K27ac signals are increased in dKO cells. **f**, RNA-seq, ChIP-seq, and ATAC-seq coverage tracks for EOMES- and BRACHYURY-repressed target genes *Sox1, Pou3f1, Neurog2, Nanog, Sox2,* and *Lefty2*. **g**, Transcription factor-binding motif enrichment within accessible chromatin in WT over dKO cells (ATAC-seq). Motifs of factors with functions during ME differentiation are enriched in WT open chromatin. p-values are indicated in g and h. **h**, Motif enrichment in ATAC-peaks of dKO compared to WT cells shows highly significant motif enrichment of factors of the pluripotency network (POU5F1, SOX2, TCF-NANOG, and KLF4) and NE programs (BRN1 (alias POU3F3) and POU3F1).

To study Tbx factor-dependent changes in chromatin accessibility, we performed Assays for Transposase-Accessible Chromatin and sequencing (ATAC-seq) and ChIP-seq for H3K27ac, a histone mark for active enhancers and transcribed regions in 5 day differentiated WT and dKO cells. We plotted ATAC sites containing EOMES- or BRACHYURY-binding peaks associated to genes downregulated in dKO cells (i.e. ME-associated sites, n=1,528) and compared signals in WT and dKO cells (Fig. 4c). Notably, dKO cells show a reduction or complete absence of chromatin accessibility at those sites, accompanied by reduced H3K27 acetylation. This observation was confirmed for several enhancers of known Tbx-activated targets such as *Mesp1, Mesp2, Cer1, Msgn1, Lhx1, Hand1,* and *Mixl1* using normalized coverage tracks of RNA-seq, ATAC-seq, and ChIP-seq data in WT and dKO cells (Fig. 4d). In contrast, accessible chromatin sites that overlap with EOMES- and BRACHYURY-binding and associate to genes upregulated in dKO (i.e. EPI- and NE-associated sites, n=743), are equally present in both WT and dKO cells (Fig. 4e). However, H3K27ac levels are increased at the same enhancer sites in dKO cells compared to WT, as expected for transcribed genes (Fig. 4e). The presence of similar levels of chromatin accessibility at pluripotency and NE enhancers in WT and dKO cells was confirmed by normalized coverage tracks of EPI genes *Nanog*, *Sox2,* and *Lefty2* and NE genes *Sox1*, *Pou3f1,* and *Neurog2* (Fig. 4f). To compare the chromatin accessibility of Tbx targets during pluripotent state and early differentiation, we analyzed ATAC-seq from mouse ESCs^40^ and EpiSCs^41^ alongside with WT and dKO ATAC-seq data. Coverage tracks show inaccessible chromatin at enhancers of ME genes in pluripotent cells, which becomes accessible in WT cells during ME differentiation and Tbx factor binding, but not in dKO cells (Supplementary Fig. 4m). In contrast, chromatin at the enhancers of pluripotency and NE genes is found in an accessible conformation already in pluripotent cells that remains largely unchanged in differentiated WT and dKO cells (Supplementary Fig. 4n). This confirms that open chromatin conformation at NE enhancers is the default state in pluripotent cells, while the accessibility of ME enhancers depends on Tbx-mediated chromatin remodeling. To compare the regulatory landscape of accessible chromatin, we performed transcription factor motif analysis of ATAC sites of WT and dKO cells and found strong enrichment of GATA, FOXA, TBX, GSC (homeobox), and MEIS1 motifs in ATAC peaks of WT cells, reflecting ME program activation (Fig. 4g; Supplementary Table 5). In contrast, ATAC sites of dKO cells are enriched for DNA-binding motifs of POU-domain homeobox (POU5F1, POU3F1), HMG (SOX2), bHLH family (BRN1), and KLF transcription factors (Fig. 4h; Supplementary Table 5), demonstrating broad differences in regulation of different lineage-specifying programs in WT and dKO cells. In conclusion, besides occupying and opening the enhancers of ME genes and acting as transcriptional activators, EOMES and BRACHYURY also bind the enhancers of EPI and NE genes that are activation-primed by accessible chromatin in pluripotent cells to transcriptionally repress them.

### EOMES and BRACHYURY repress EPI and NE genes directly and indirectly by ME gene networks

We next analyzed the repressive functions of EOMES and BRACHYURY in more detail by generating constructs encoding VP16 transactivation- or Engrailed repression-domains (EnR) C-terminally fused to the Tbx DNA-binding domain of truncated EOMES or BRACHYURY, then acting as dominantly activating or repressing fusion proteins^42, 43^. A N- and C-terminally truncated version of EOMES containing only the DNA-binding domain (EoTBX) was used to test if DNA-binding by itself can act repressively (Fig. 5a). Resulting constructs (EoVP16, BraVP16, EoEnR, BraEnR, and EoTBX) were introduced into the dox-inducible locus of dKO cells while full-length cDNAs (EoFL, BraFL) were used as controls. ChIP-seq analysis of cells expressing EoGFP, EoFL, and EoVP16 indicated highly similar chromatin binding of the different fusion proteins as the FL EOMES protein (Supplementary Fig. 5a). We performed transcriptional profiling of day 5 differentiating dKO cells after dox-induced expression of constructs from day 3 to day 5 (Supplementary Fig. 4d) and analyzed the expression of deregulated genes in dKO cells (Fig. 5c-f; Supplementary Table 6). EoVP16 and BraVP16 constructs effectively rescued expression of ME genes similar to control EoFL (n=260 overlapping genes) and BraFL (n=150 overlapping genes) constructs, including known EOMES and BRACHYURY target genes (Fig. 5c,d,i; Supplementary Fig. 5b,d,e; Supplementary Table 6). Remarkably, EoEnR and BraEnR repressor constructs, as well as EoFL (n=75 overlapping genes) and BraFL (n=72 overlapping genes) strongly reduced levels of EPI and NE genes in dKO cells (Fig. 5e,f,j; Supplementary Fig. 5c,f,g; Supplementary Table 6), supporting direct repression by Tbx factors. The repression of pluripotency and neural fate in dKO cells expressing the different rescue constructs was confirmed by IF staining for the pluripotency marker NANOG and the neural marker TUJ1. Both markers are abundantly present in dKO cells, while numbers of NANOG and TUJ1 positive cells are drastically reduced following the expression of the FL and VP16 constructs (Fig. 5g,h). Reduced numbers of NANOG positive cells are observed following EoEnR, but not after BraEnR expression. In contrast, BraEnR expression reduces the number of TUJ1 positive neurons, while EoEnR has little effect (Fig. 5g,h). In accordance, mRNA levels of pluripotency genes *Nanog*, *Pou5f1*, *Lefty2*, *Wnt8a* and others are predominantly reduced by EoEnR (Fig. 5e,j; Supplementary Fig. 5c,f), whereas BraEnR reduces NE gene expression, such as *Sox1*, *Sox2, Sox3,* and *Nkx6-2* (Fig. 5f,j; Supplementary Fig. 5c,g) showing certain specificity in repressive functions of *Brachyury* and *Eomes*.

**Fig. 5:**
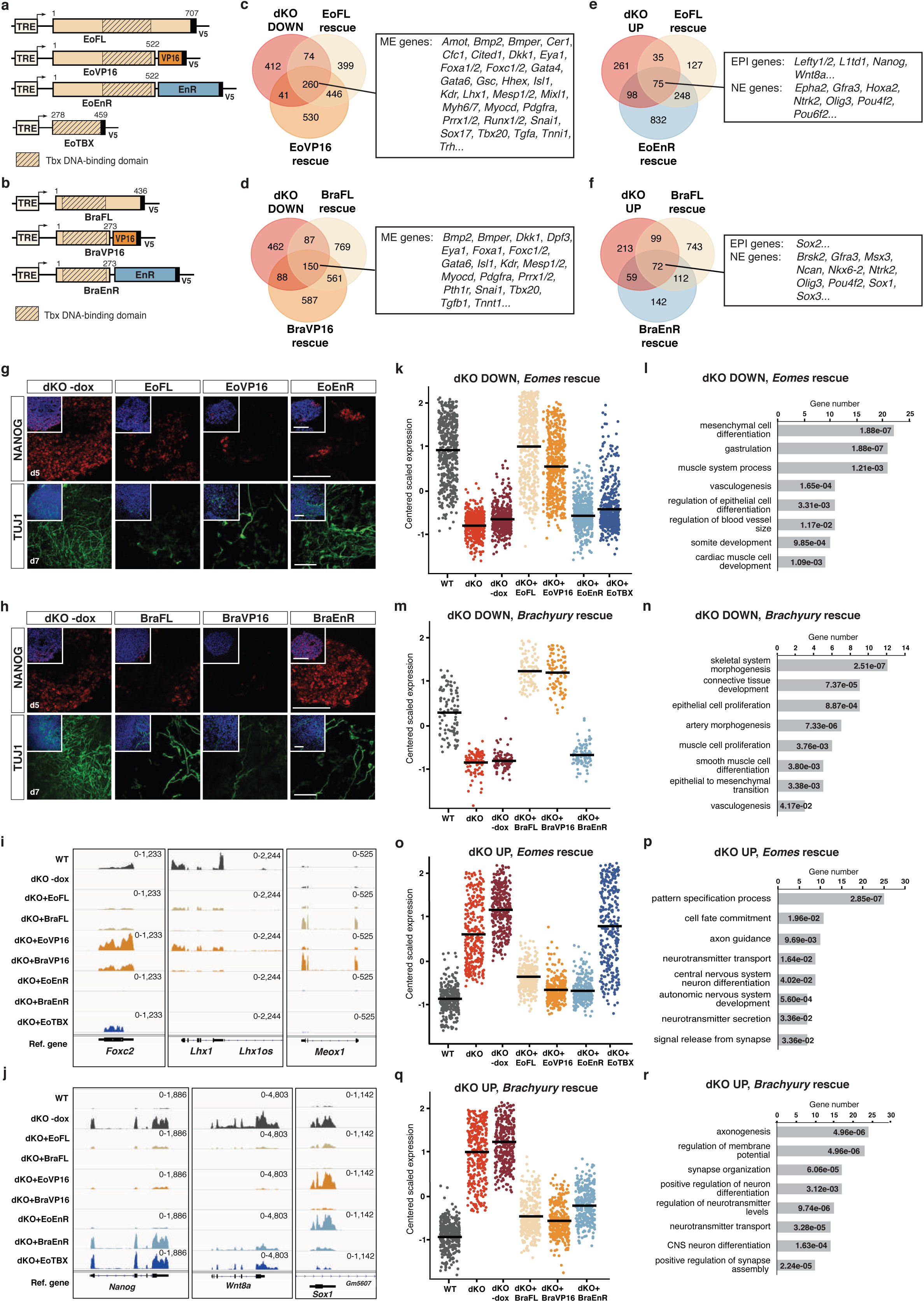
Tbx factors repress EPI and NE genes directly and indirectly by activation of downstream ME target genes. **a,b**, Schematic of *Eomes* and *Brachyury* FL (full-length), VP16 (activator), EnR (repressor), and EoTBX constructs inserted into the doxycycline (dox)-inducible locus (TRE) of dKO ESCs. **c,d**, Overlap of genes downregulated in dKO cells that are rescued by (c) EoFL and EoVP16, and (d) BraFL and BraVP16 when analyzed by RNA-seq. Adjusted p-value≤0.05, log_2_(FC) of +/-1.5 was used for rescue constructs in c-f. Representative genes of ME differentiation are indicated. **e,f**, Overlap of genes that are upregulated in dKO cells and reduced upon expression of (e) EoFL and EoEnR, and (f) BraFL and BraEnR. Representative genes indicative for EPI and NE are boxed. **g**, IF staining showing numbers of NANOG positive cells at day 5 of differentiation and TUJ1 positive cells at day 7 of differentiation after induced expression of EoFL, EoVP16, and EoEnR. Scale bars 100 μm in g and h. **h**, IF staining for NANOG and TUJ1 in BraFL, BraVP16, and BraEnR expressing cells on day 5 and 7 of differentiation, respectively. **i**, RNA-seq coverage tracks following induced expression of rescue constructs shown in a and b for indicated mesoderm genes (see also Supplementary Fig. 5b). Counts normalized to RPKM are indicated in i and j. **j**, RNA-seq coverage tracks for EPI markers (*Nanog* and *Wnt8a*) and the NE marker *Sox1* in rescue experiments, showing direct repression of EPI genes by *Eomes* and of NE markers by *Brachyury* EnR constructs (see also Supplementary Fig. 5c). **k,m**, Plots depicting centered scaled counts of downregulated genes in dKO (red) and uninduced dKO-dox cells (dark red), and after induction of indicated Eomes- (k) or Brachyury- (m) rescue constructs. Black line shows mean counts within each sample in k,m,o, and q. Gene expression is rescued to WT levels by FL and VP16-activator constructs, but not by EnR-repressor and EoTBX. **l,n**, GO-term analyses of genes depicted in k and m shows enrichment of terms associated with ME development. The p-values for each term are indicated in l-r. **o,q**, Plots depicting centered scaled counts of upregulated genes in dKO (red), and uninduced dKO-dox cells (dark red), and count values of the same genes following the expression of Eomes- (o) or Brachyury- (q) rescue constructs. Expression of selected genes is reduced by all three types of rescue constructs (FL, VP16, and EnR), but not by EoTBX. **p,r**, GO-term analyses of genes depicted in o and q showing enrichment of terms associated with neural development.

To further analyze the transcriptional responses to the transactivator and repressor constructs, we performed unsupervised clustering of differentially expressed genes of dox treated compared to untreated cells (Fig. 5k,m,o,q; Supplementary Table 7) followed by GO-term analysis (Fig. 5l,n,p,r; Supplementary Table 7). This analysis demonstrated that Tbx-induced ME genes are activated by the FL and VP16 constructs, but not by EnR constructs (Fig. 5k-n). Nonetheless, EoEnR and BraEnR induction effectively reduced EPI and NE markers to the levels of WT cells (Fig. 5o-r), supporting a direct mode of repression. Expression of only the TBX domain was not sufficient to activate or repress target genes (Fig. 5 k,o). Surprisingly, genes of the EPI and NE programs that are directly repressed by EnR constructs additionally show downregulation following the induction of EoVP16 and BraVP16 (Fig. 5o,q; Supplementary Fig. 5f,g), as shown in previous IF staining (Fig. 5g,h). Since such regulation does not comply with a simple, linear mode of direct repression of EPI and NE genes by Tbx factors alone, we hypothesized that EPI and NE target genes are subject to additional repression by Tbx-induced factors that are most likely the components of downstream ME gene programs.

### ME-specifying factors repress pluripotency and NE programs in the absence of Tbx factors

Next, to explore if EPI and NE genes are additionally repressed by Tbx-induced early ME-specifying factors, we generated dKO cells that allow for dox-inducible expression of the mesoderm (Mes) transcription factors *Mesp1* and *Msgn1* and definitive endoderm (DE) factors *Mixl1* and *Foxa2* (Fig. 6a). These represent known directly regulated targets genes of *Eomes* or *Brachyury* that are expressed in the earliest populations of Mes and DE progenitors (Fig. 1c; Supplementary Fig. 5a,b; Supplementary Fig. 6a). *Six2*, a marker for head and heart mesenchyme progenitor populations from E8.5 was chosen as a late acting mesenchymal regulator^44^. Resulting ESCs were differentiated according to the previous protocol (Supplementary Fig. 4d) and analyzed by RNA-seq. PCA shows close clustering of *Mesp1* and *Msgn1*, and of *Mixl1* and *Foxa2* expressing cells in separate groups between WT and dKO cells, indicating partial rescues of the dKO phenotype, while *Six2* expressing cells remained closely clustered to the dKO cells (Fig. 6b). Transcriptome analysis indicates activation of similar sets of early Mes-specific genes following *Mesp1* and *Msgn1* expression (Fig. 6c; Supplementary Fig. 6b,c; Supplementary Table 8), and both Mes and DE genes following *Mixl1* and *Foxa2* expression (Fig. 6d; Supplementary Figure 6d,e; Supplementary Table 8). *Six2* failed to induce early Mes marker genes, but among others induced key factors for early metanephric mesenchyme, such as *Pax2* and *Sall1* (Fig. 6c,d; Supplementary Fig. 6f). Strikingly, all four factors, but not *Six2* effectively reduced expression of the EPI and NE markers in dKO cells to levels in WT cells (Fig. 6e-h; Supplementary Table 8). The repression of the pluripotency factor OCT4 and neuronal markers (SOX1 and TUJ1) was confirmed by IF staining, showing strongly reduced numbers of positive cells in induced compared to uninduced dKO cells (Fig. 6i).

**Fig. 6:**
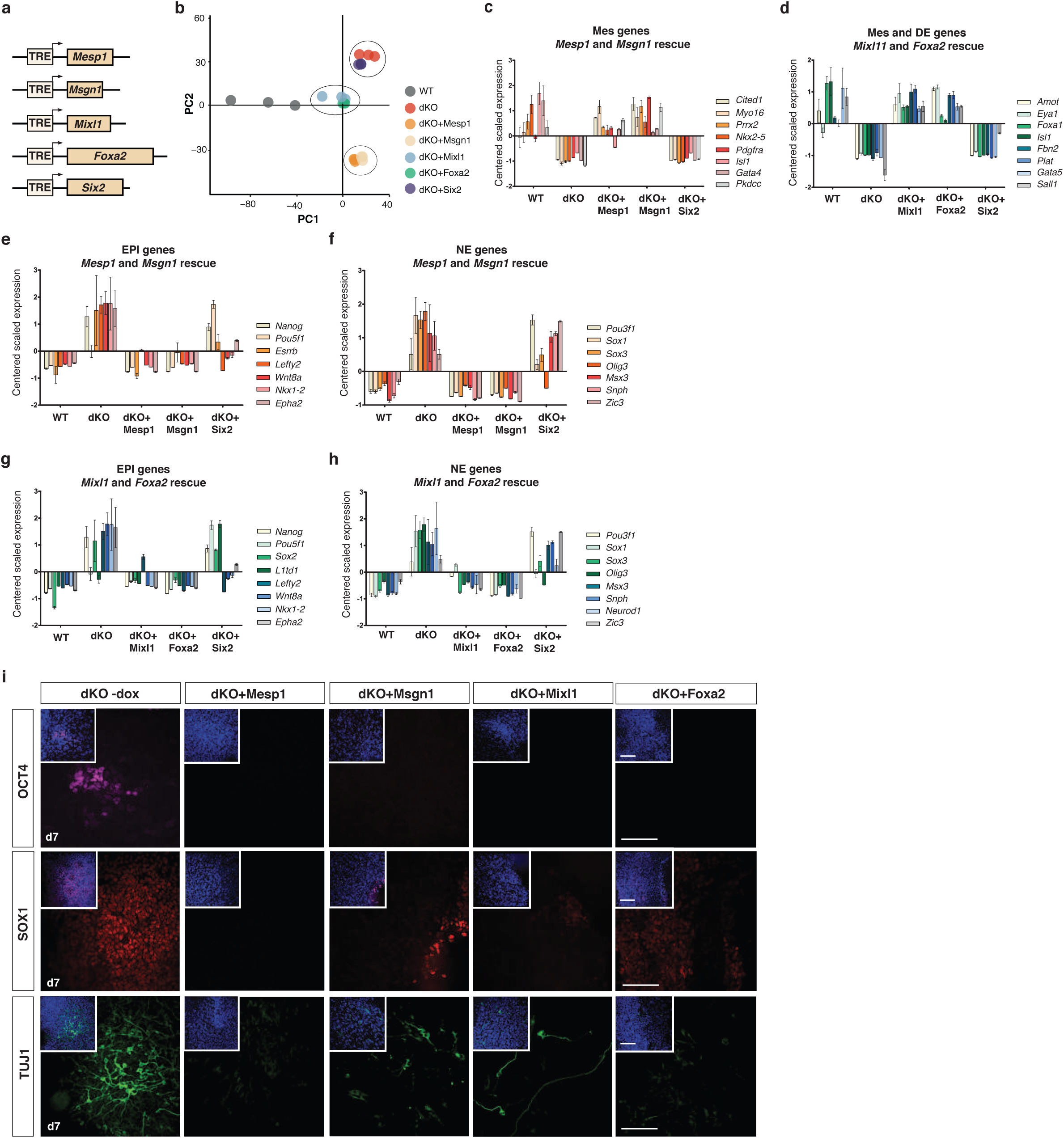
Mesoderm and endoderm transcription factors downstream of Tbx genes act as repressors of pluripotency and neuroectoderm programs. **a**, Schematic of cDNA constructs of mesoderm (Mes; *Mesp1, Msgn1*), definitive endoderm (DE; *Mixl1, Foxa2*) and mesenchymal (*Six2*) transcription factors introduced into the dox-inducible locus (TRE) of dKO cells. **b**, Principal component (PC) analysis of RNA-seq data after induced expression of indicated factors showing a partial rescue of the dKO phenotype and clustering proximity of dKO+Mesp1 and dKO+Msgn1, as well as dKO+Mixl1 and dKO+Foxa2 cells, while dKO+Six2 cells remain closely clustered to dKO cells. **c**, Rescue of Mes genes by induced expression of *Mesp1* or *Msgn1* in differentiated dKO cells. Bars represent centered scaled counts from triplicate RNA-seq experiments and error bars indicate SEM in c-h. **d**, Rescue of Mes and DE genes by induced expression of *Mixl1* or *Foxa2*. **e-h**, Expression levels of EPI (e,g) and NE genes (f,h) indicated by centered scaled counts are reduced to WT levels after induced expression of *Mesp1* and *Msgn1* (e,f) and *Mixl1* and *Foxa2* (g,h). **i**, IF staining for pluripotency marker OCT4 and neural markers SOX1 and TUJ1 at day 7 of EB differentiation showing reduced number of stained cells after induced expression of ME transcription factors *Mesp1, Msgn1, Mixl1, or Foxa2.* Scale bars 100 μm.

We conclude that pluripotency and NE programs are subjected to repression in ME-forming cells at several levels: initial direct repression by Tbx factors, and subsequent repression by their downstream targets after initiation of ME-specific transcriptional programs, ensuring proper lineage specification by the continuous suppression of competing programs.

## Discussion

In this study we address the control mechanisms of pluripotency exit and germ layer specification that lead to the segregation of the NE and ME lineages. We demonstrate essential functions of the Tbx factors *Eomes* and *Brachyury* for the activation of all ME programs, and reveal novel functions as direct repressors of key pluripotency and NE genes (Fig. 7a,b). This regulation involves chromatin remodeling on ME target gene enhancers during lineage commitment, as summarized in a comprehensive model (Fig. 7c).

**Fig. 7:**
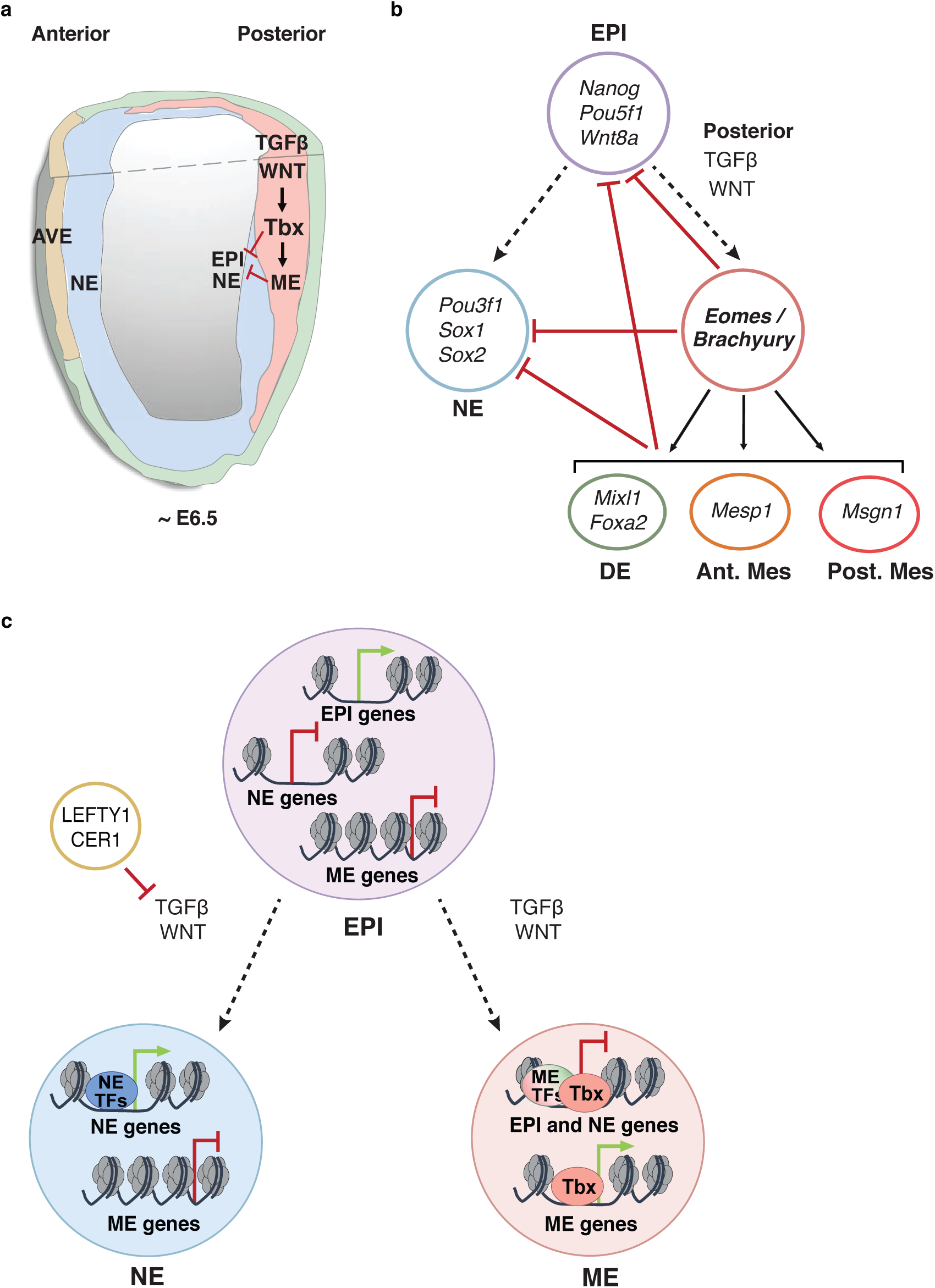
Proposed control mechanisms of pluripotency exit and cell lineage segregation by Tbx factors. **a**, Schematic of a ∼E6.5 embryo showing that rising levels of TGFβ and WNT signals induce Tbx factors *Eomes* and *Brachyury* in pluripotent epiblast (EPI) cells at the posterior side. Tbx factors induce mesoderm and definitive endoderm (ME) formation and repress primed pluripotency of the epiblast (EPI) and neuroectoderm (NE) fate. NE is specified in the anterior EPI that is shielded from TGFβ and WNT signals by secreted antagonists LEFTY1 and CER1 from the anterior visceral endoderm (AVE). **b**, Tbx factors activate the transcriptional programs for specification of anterior mesoderm (Ant. Mes), posterior mesoderm (Post. Mes) and definitive endoderm (DE). Concomitantly, Tbx factors downregulate primed pluripotency by direct repression of core regulators (*Nanog, Pou5f1, Wnt8a*) and indirectly via induced ME transcription factors (TFs) (*Mesp1, Msgn1, Mixl1, Foxa2*). NE genes, such as *Pou3f1, Sox1,* and *Sox2* are similarly repressed by Tbx and downstream ME TFs. **c**, Schematic of chromatin accessibility changes during germ layer specification. In EPI cells the NE enhancers are accessible and activation primed. ME enhancers are in closed chromatin state and only become accessible in response to Tbx and ME TFs binding, that repress EPI and NE gene transcription in ME-forming cells. NE genes initiate transcription in NE-forming cells in the absence of repression by Tbx and other ME TFs. Dashed lines indicate differentiation paths; black arrows indicate activation; red lines with bars indicate repression.

Previous reports have shown that *Eomes* and *Brachyury* are indispensable for the initiation and specification of distinct ME cell lineages^7, 8, 17, 33^ and are found from the earliest timepoints of gastrulation onset in the mouse embryo^1, 45^ and in differentiating human^17^ and mouse ESCs (this study). Despite the large overlap in chromatin occupancy at regulatory sites of target genes^14–16^, *Eomes* and *Brachyury* fulfill different, non-overlapping, potentially competing functions during ME specification^46^ as also reflected in the different loss-of-function phenotypes^7, 10, 11^. Current data show that the genetic deletion of *Eomes* results in the increased expression of *Brachyury* (Supplementary Fig. 1c) and its target genes, and *vice versa* (Fig. 1c,e). This suggests mutual feedback regulation that is currently not understood in detail and requires further studies. However, only the simultaneous deletion of both Tbx factors reveals their fundamental role for the activation of the gene programs of all ME lineages, resulting in the gross reduction of ME enhancer accessibility in the absence of Tbx factors^47, 48^.

In addition, we identified novel functions of *Eomes* and *Brachyury* as transcriptional repressors of pluripotency genes resulting in a gene signature resembling the pluripotent state of the pregastrulation epiblast in dKO cells^49^. Here, *Eomes* and *Brachyury* show rather synergistic functions for the repression of pluripotency programs. Of note, binding of EOMES to regulatory sites of core pluripotency factors *POU5F1, NANOG,* and *SOX2* in human ESCs differentiation had previously been reported^17^. However, the knockdown of *EOMES* in these experiments showed only minor effects on expression levels of pluripotency genes, owing to redundant functions with *BRACHYURY*, as suggested by our data. Despite the absence of ME program activation dKO cells eventually downregulate pluripotency and undergo differentiation to NE cell types when cells are removed from pluripotency maintaining conditions. This suggests that different modes of pluripotency exit are employed during ME and NE differentiation.

Previous studies had demonstrated that TGFβ and WNT signals are required to restrict NE lineage specification^22^, yet the exact molecular mechanism remained elusive. We demonstrate that *Eomes* and *Brachyury* are the crucial effectors of TGFβ and WNT signaling that mediate NE fate repression, since dKO cells differentiate into neurons in the presence of high doses of ActA. In addition to direct repression by Tbx factors, Tbx-induced transcriptional regulators of ME-specifying programs, such as *Mesp1*, *Msgn1*, *Mixl1*, and *Foxa2*, and likely also others efficiently repress pluripotency and NE lineage signatures of dKO cells, as previously shown for *Mesp1*^50^. This represents a remarkable example of synergistically acting repression by cascades of hierarchically-acting ME transcription factors that ensure robust NE and pluripotency program silencing. Such degree of redundancy is advised when expression levels of highly dynamic transcription factors as *Eomes* and *Brachyury* are rapidly downregulated in the course of specification, to ensure the continuous repression of the opposing lineage programs and maintenance of the acquired ME cell fate.

The repression of neuronal differentiation by Tbx factors was previously reported in different contexts, including the bipotential axial progenitors of late gastrulating mouse embryos (>E8.5)^51^. Axial progenitors are characterized by the dual expression of *Brachyury* and *Sox2* that antagonistically control mesodermal or neural fate, respectively^52, 53^. Moreover, the genetic deletion of *Tbx6*, leads to the excessive generation of neural tissues at the expense of paraxial mesoderm^54, 55^. Studies in *Xenopus* had shown that the combined knockdown of the three Tbx factors *eomes, vegT,* and *xbra/xbra3* causes an oversized neural tube and a failure to form mesoderm caudal to head region^16^. Interestingly, *Eomes* (alias *Tbr2*) is also implicated at later developmental stages (>E10.0) in cortical brain development^56^. Here, *Eomes* contributes to the generation of neurogenic basal progenitors, by suppressing key neuronal stemness factors of radial glia, such as *Pax6* and *Insm1*^57^. However, these studies addressed fate decisions at later stages of development, and did not investigate if the Tbx factors act as direct repressors or via induced downstream factors.

Interestingly, current experiments using the EnR repressor constructs in dKO cells suggested certain specificity for the direct repression of pluripotency and NE genes by *Eomes* and *Brachyury*. We found that EOMES represses pluripotency, while BRACHYURY predominantly inhibits NE gene expression. This is in accordance with the embryonic functions of *Eomes* in the early epiblast^8, 17^ and *Brachyury* at later embryonic stages e.g. in bipotential neuromesodermal progenitors^51–53^.

Despite the wealth of knowledge on transcriptional regulation of lineage specification by *Eomes* and *Brachyury*, the exact molecular mechanisms are just starting to be explored^58–62^. Previous reports have suggested that pluripotent cells have globally open chromatin that becomes progressively restricted during differentiation^63^. However, a recent report suggested global differences in chromatin accessibility and methylation patterns between NE and ME gene enhancers in the early pluripotent epiblast^64^. Here, we demonstrate that Tbx factors are indispensable for the establishment of accessible chromatin at ME enhancers. This suggests that EOMES and BRACHYURY bind to otherwise inaccessible chromatin and recruit chromatin remodeling complexes to target genes establishing competence for their activation, similar to “pioneering factors”^65^. This is further supported by a recent study showing that in addition to binding to free DNA, Tbx, among other transcription factors, also interact with nucleosome-bound DNA, possibly binding DNA gyre of neighboring nucleosomes as dimers^66^. In contrast to ME enhancers, the enhancers of pluripotency and NE genes reside in an open chromatin state in pluripotent cells and the expression of these genes is actively repressed during ME specification. We did not detect significant differences in chromatin accessibility of NE enhancers occupied by Tbx factors between WT and dKO cells, implying the existence of other mechanisms for Tbx-mediated repression of pluripotency and NE genes^67^. The expression of the Tbx DNA-binding domain of *Eomes* had no major effect on gene activation, or repression in dKO cells, suggesting that other mechanisms than mere DNA-binding are required for activation, or repression of target genes. A previous report described the interaction of another Tbx factor, TBX5, with the NuRD (Nucleosome Remodeling and Deacetylase) complex during cardiac development for the repression of target genes^68^. More thorough analyses of interacting proteins within nuclear complexes will reveal the various mechanisms underlying transcriptional control and chromatin remodeling by EOMES and BRACHYURY.

The remarkable differences of chromatin accessibility at regulatory sites of NE and ME genes present in pluripotent cells might explain the differentiation bias of pluripotent cells towards NE in the absence of ME-inducing signals. Future studies should address the mechanisms that prevent premature NE gene activation in pluripotent cells, as well as the mechanisms for their timely activation, e.g. by transcriptional regulators that promote NE fate when pluripotent cells initiate differentiation, such as *Sox2* and *Pou3f1*^27^. It will also be interesting to learn if preexisting, asymmetric lineage priming in combination with repressively-acting transcription factors represents a common mode of binary cell fate decisions during differentiation of stem and progenitor cells.

## Acknowledgements

We thank T. Bass and C. Domisch for excellent technical assistance; B.G. Herrmann for providing *T*^2J^ mice and M. Kyba for the A2lox.Cre ESC line. We are thankful to the Freiburg Galaxy Team, especially B. Grüning (Bioinformatics, University of Freiburg, Germany) and computing support by the state of Baden-Württemberg through bwHPC, University of Tübingen. We thank C. Schwan for imaging support; S. Prekovic for valuable advice concerning ChIP; S. Nothjunge and R. Gilsbach for technical advice and comments on the data; M. Timmers for advice on GFP-fusion constructs; C. Hill and T. Gaarenstroom for the pGL2-6xARE-Lux plasmid; G. Schmidt and S. Kowarschick for help with luciferase assays, D. Onichtchouk, M.A. Morgan and P. Walentek for critical reading of the manuscript and discussions, and the staff of the Life Imaging Centre (Center for Biological Systems Analysis, Albert-Ludwigs-University Freiburg) for help with confocal microscopy. We acknowledge the Genomics Core Facility at the EMBL (Heidelberg, Germany) for sequencing. This study was supported by the German Research Foundation (DFG) through the Emmy Noether- and Heisenberg-Programs (AR 732/1-1/2/3, AR 732/3-1), project grant (AR 732/2-1,) project B07 of SFB 1140 (project ID 246781735), project A03 of SFB 850 (project ID 89986987), and Germany’s Excellence Strategy (CIBSS – EXC-2189 – Project ID 390939984) to S.J.A.; TRR 152 (project ID 239283807) project P03, and Germany’s Excellence Strategy (CIBSS – EXC-2189 – Project ID 390939984) to M.K.; and by the DFG SFB 992 (project ID 192904750) project B03 to L.H.

## Author Contribution

J.T., M.P., C.M.S., S-L.M., M.B., S.P., and S.J.A. performed experiments. J.T. and S.J.A. planned and analyzed experiments. J.T. and G-J.K. performed bioinformatics data analysis. A.H. and M.K. designed TALENs. L.H. analyzed the data and edited the manuscript. J.T. and S.J.A. prepared figures, wrote and edited the manuscript with input from all authors. S.J.A. conceived the study.

---

**Supplementary Fig. 1.**
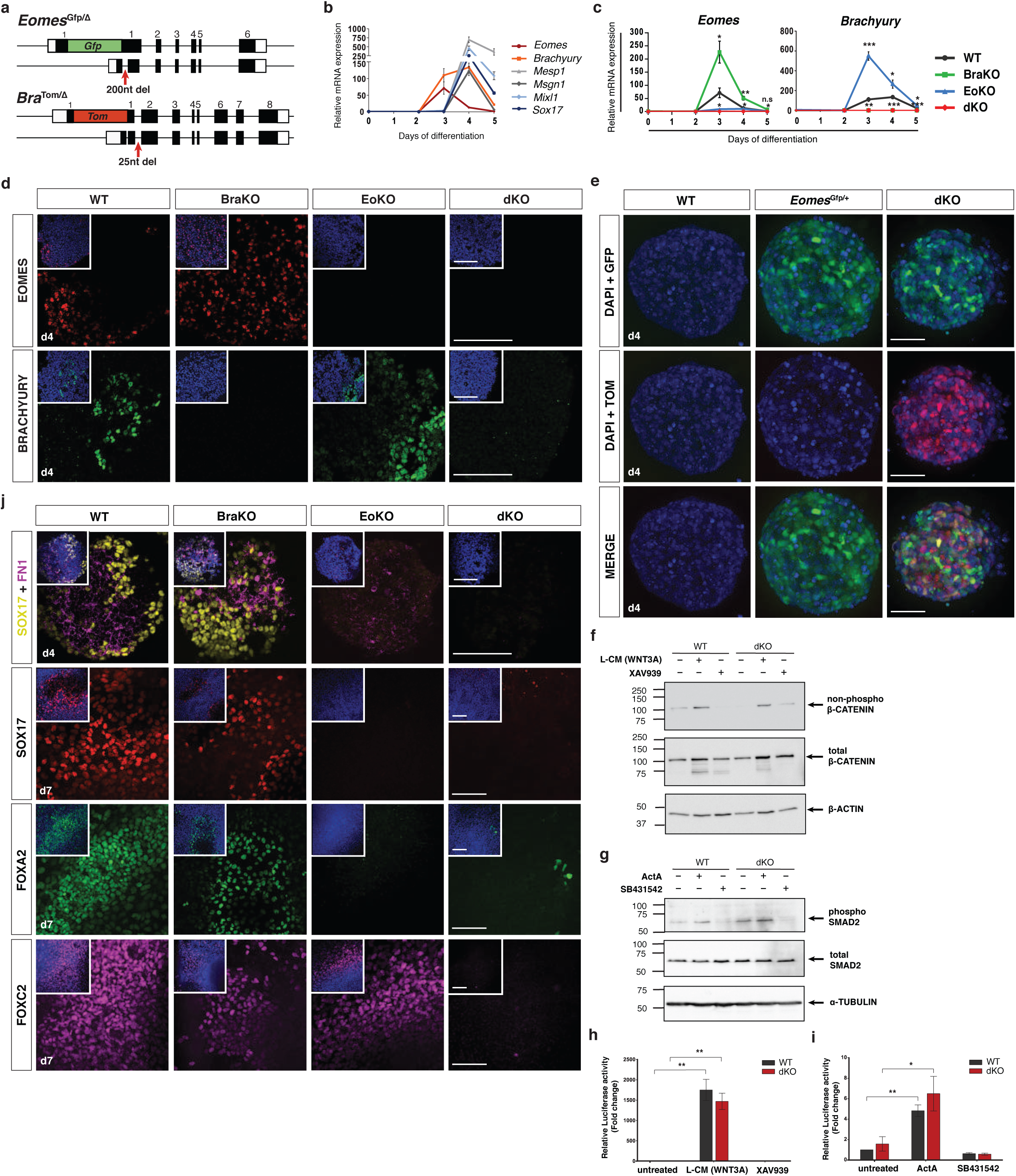
Characterization of dKO cells for EOMES and BRACHYURY expression, reporter activation, and differentiation potential. **a**, Schematic of *Eomes*^Gfp/Δ^ (EoKO) and *Bra*^Tom/Δ^ (BraKO) alleles. Fluorescence reporters Gfp and Tom, including polyA signals were inserted into the start codons of one allele of *Eomes* and *Brachyury*, respectively, and the second allele is functionally disrupted using TALENs to generate short out-of-frame deletions within the first exon. **b**, Relative mRNA expression of mesoderm and endoderm (ME) genes alongside with *Eomes* and *Brachyury* expression over 5 days of differentiation of WT cells. Error bars represent SEM. **c**, Expression levels of *Eomes* and *Brachyury* transcripts during the 5 days of differentiation of WT, BraKO, EoKO, and dKO cells measured by qRT-PCR. Error bars indicate SEM; p-values for differences of mean expression between WT and dKO samples were calculated by Student’s t-test. n.s:p>0.05; *:p≤0.05; **:p≤0.01; ***: p≤0.001; ****: p≤0.0001. **d**, Immunofluorescence staining at day 4 of differentiation demonstrating the absence of EOMES and BRACHYURY in respective loss-of-function cell lines. Scale bars 100 μm. **e**, *Eomes*^Gfp^ and *Bra*^Tom^ reporter activation in *Eomes*^Gfp/+^ and dKO EBs at day 4 of differentiation. Maximum intensity projection of confocal z-stacks is shown. Scale bars 100 μm. **f**, Protein levels of non-phosphorylated (active) β-CATENIN and total β-CATENIN in WT and dKO cells showing responsiveness to WNT stimulation. β-ACTIN served as the loading control. **g**, Protein levels of phosphorylated SMAD2 and total SMAD2 in WT and dKO cells showing responsiveness to ActA stimulation. ɑ-TUBULIN served as the loading control. **h**, Super 8x TOPflash luciferase reporter assay demonstrating responsiveness of WT and dKO cells to WNT stimulation when treated with WNT3A L-cell conditioned medium (CM), but not when untreated or inhibited with XAV939. Error bars indicate SEM. p-values for differences of mean expression between treated and untreated samples were calculated by Student’s t-test. *:p≤0.05; **:p≤ 0.01 in h and i. **i**, 6xARE Luciferase reporter assay demonstrating responsiveness of WT and dKO cells to TGFβ/Nodal signalling when treated with ActA, but not when untreated or inhibited with SB431542. **j**, Immunofluorescence staining for SOX17, FIBRONECTIN1 (FN1), FOXA2, and FOXC2 proteins in plated EBs at day 4 and 7 of differentiation showing the absence of endoderm (SOX17 and FOXA2) and mesoderm (FN1 and FOXC2) markers. Scale bars 100 μm.

**Supplementary Fig. 2:**
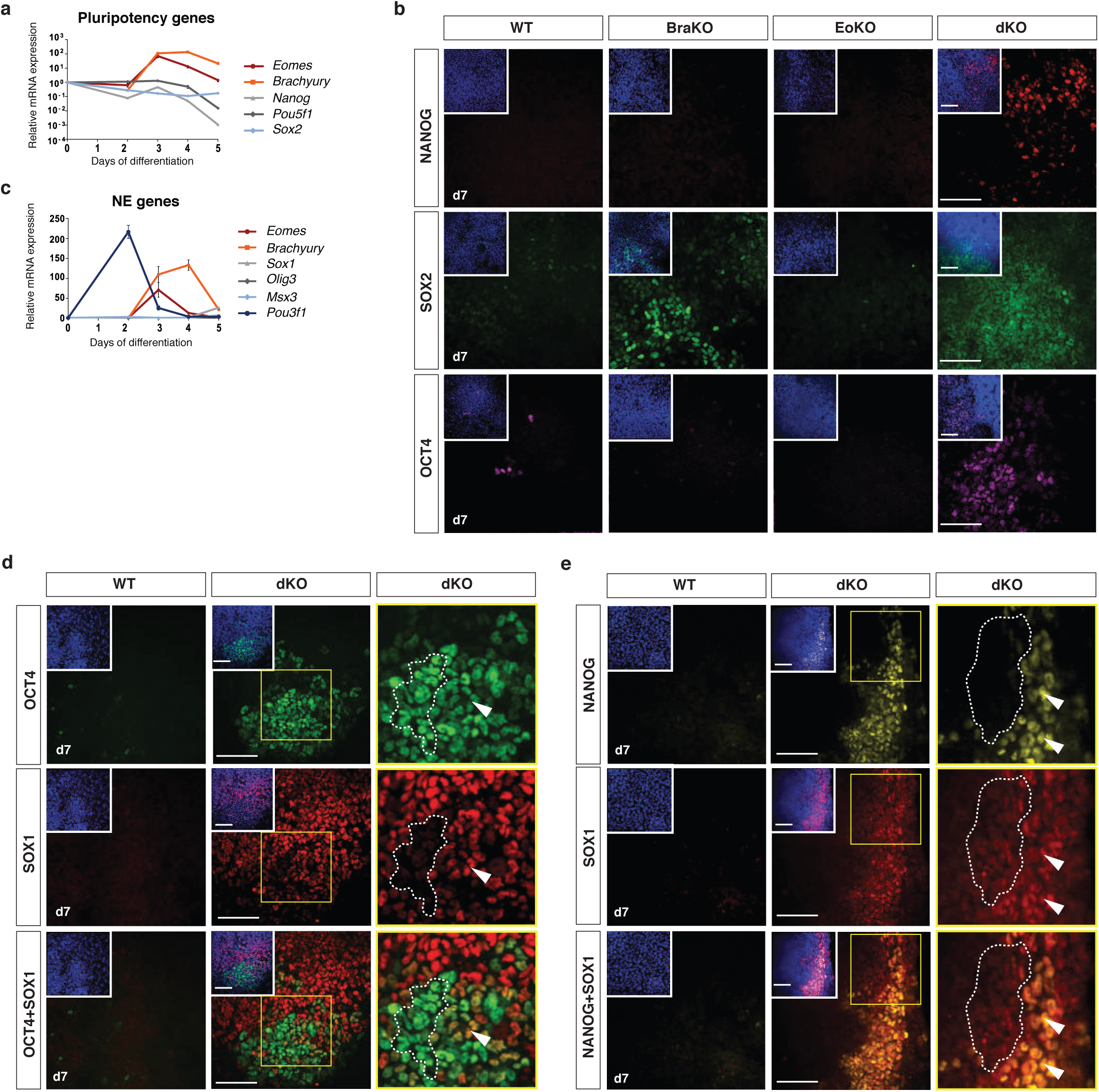
*Eomes*- and *Brachyury*-deficient cells retain pluripotency and express neuroectoderm markers during differentiation. **a**, Relative mRNA expression of pluripotency genes alongside with *Eomes* and *Brachyury* expression over 5 days of differentiation of WT cells. Error bars represent SEM. **b**, Immunofluorescence staining for NANOG, SOX2, and OCT4 of plated EBs at day 7 of differentiation showing maintained expression of pluripotency markers in dKO cells, that are lost during differentiation of WT, BraKO, and EoKO cells. Scale bars 100 μm. **c**, Relative mRNA expression of neuroectoderm (NE) genes alongside with *Eomes* and *Brachyury* expression over 5 days of differentiation of WT cells. Error bars represent SEM. **d**, Co-immunofluorescence staining in dKO EBs at day 7 of differentiation showing OCT4 and SOX1 co-expression in a small proportion of cells (arrowheads), few cells express only OCT4 (dashed line), and most cells express only SOX1. Yellow rectangles are shown at higher magnification to the right. Scale bars 100 μm. **e**, Co-immunofluorescence staining in dKO EBs at day 7 of differentiation showing NANOG and SOX1 co-expression in few cells indicated by arrowheads. Most cells show only SOX1 staining (dashed line). Single NANOG positive cells are not detected. Yellow rectangles are shown at higher magnification to the right. Scale bars 100 μm.

**Supplementary Fig. 3:**
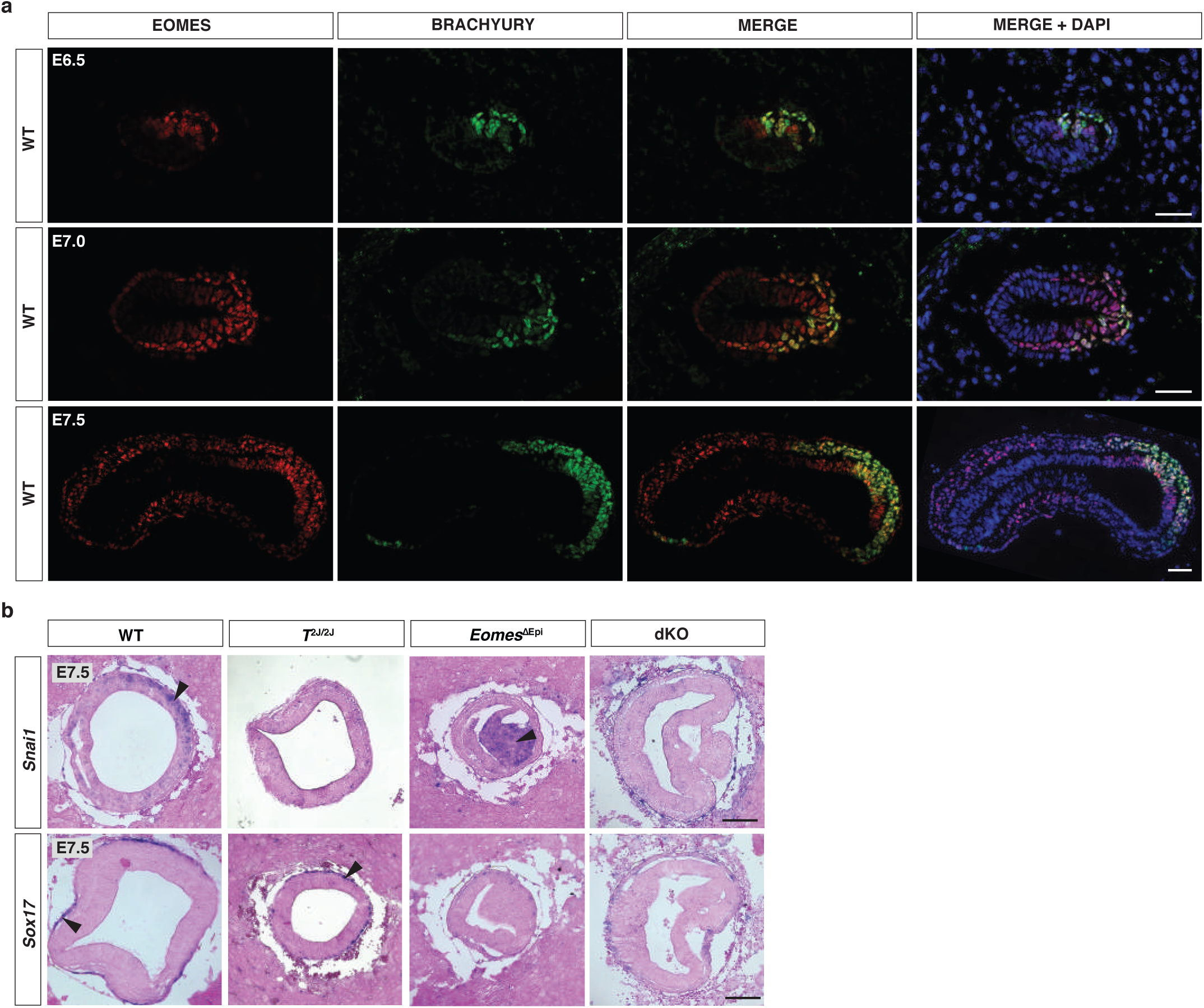
EOMES and BRACHYURY co-expression in the posterior epiblast and nascent mesoderm during gastrulation onset. **a**, Immunofluorescence staining of transverse sections of E6.5, E7.0, and E7.5 WT embryos showing EOMES and BRACHYURY co-expression in posterior-proximal epiblast and nascent mesoderm at E6.5. At E7.0 and E7.5 EOMES expression extends more anteriorly in the epiblast than BRACHYURY. Double positive cells are found in the epiblast and in nascent mesoderm. Scale bars 50 μm. **b**, mRNA *in situ* hybridization analysis of transversal sections of E7.5 embryos of indicated genotypes for *Snai1* and *Sox17* showing absence of both markers in the dKO embryos. Arrowheads indicate sites of staining for *Snai1* and *Sox17* expression. Scale bars 100 μm.

**Supplementary Fig. 4:**
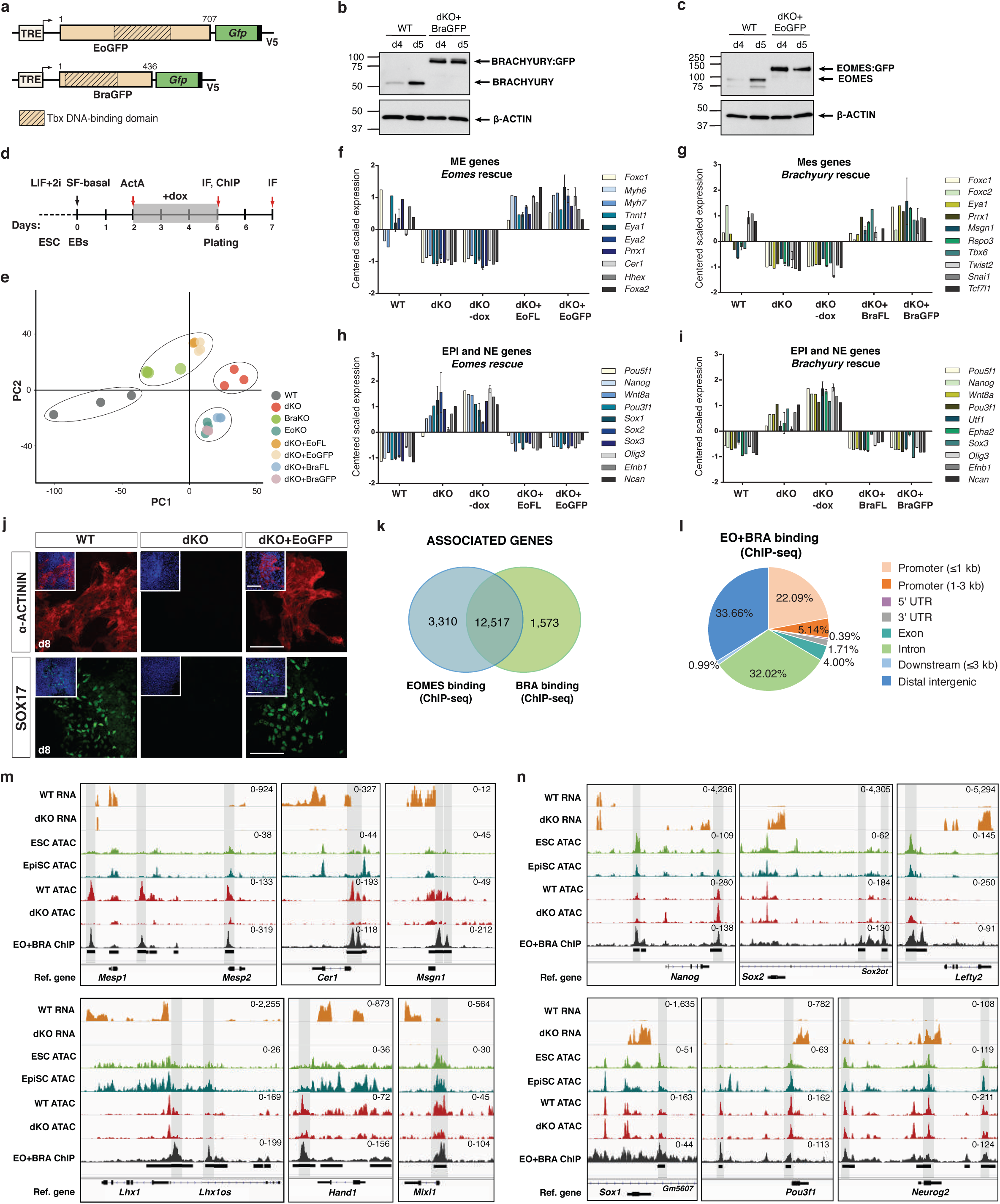
EOMES and BRACHYURY bind to regulatory regions of ME genes, as well as EPI and NE genes, and impact on chromatin accessibility. **a**, Schematic of EoGFP and BraGFP constructs used for ChIP-seq experiments. GFP was fused to the C-terminus of full-length *Eomes* or *Brachyury* and introduced into the dox-inducible locus (TRE) of dKO ESCs. **b,c**, Protein leves of BRACHYURY (b) and EOMES (c) in WT cells compared to dKO+BraGFP and dKO+EoGFP cells after 4 and 5 days of differentiation. β-ACTIN served as the loading control. **d**, Differentiation protocol of dKO+EoGFP and dKO+BraGFP cells to ME by administration of dox from day 2 to day 5 of differentiation. **e**, Principal component (PC) analysis of RNA-seq expression data of indicated cell lines at day 5 of differentiation. dKO+EoGFP and dKO+EoFL cells cluster more closely to BraKO cells, and dKO+BraGFP and dKO+BraFL cluster more closely to EoKO cells, indicating functional rescues by GFP fusion constructs. **f-i**, Expression levels indicated by centered scaled counts of mesoderm and endoderm (ME) and pluripotency and neuroectoderm (EPI and NE) genes that are rescued after induced expression of *Eomes-* (f, h) or *Brachyury-* (g, i) FL and GFP constructs. Bars represent centered and scaled mRNA expression levels obtained by triplicate experiments of RNA-seq and error bars indicate SEM. **j**, Immunofluorescence staining for α-ACTININ and SOX17 of plated EBs at day 8 of differentiation indicating cardiomyoyte and endoderm differentation of WT and dKO+EoGFP cells, but not of dKO cells. Scale bars 100 μm. **k**, Overlap of genes associated to regions bound by EOMES or BRACHYURY showing that the vast majority of genes contain ChIP-seq peaks for both Tbx factors. **l**, Genomic distribution of EOMES- and BRACHYURY-bound sites (ChIP-seq) showing predominant binding to regions in the proximity to gene bodies. **m**, RNA-seq, ATAC-seq, and ChIP-seq coverage tracks of differentiated WT and dKO cells, ESCs^40^ and EpiSCs^41^ at loci of proposed Tbx-activated target genes. ATAC peaks at regulatory sites that were not present in pluripotent cells (ESCs and EpiSCs) are established during differentiation to ME and are bound by EOMES and BRACHYURY. Counts normalized to RPKM are indicated in the right corner in m and n. **n**, RNA-seq, ATAC-seq, and ChIP-seq coverage tracks of potentially Tbx-repressed pluripotency (*Nanog*, *Sox2*, *Lefty2*) and NE (*Sox1*, *Pou3f1*, *Neurog2*) target genes showing that chromatin is already accessible in pluripotent cells (ESCs and EpiSCs).

**Supplementary Fig. 5:**
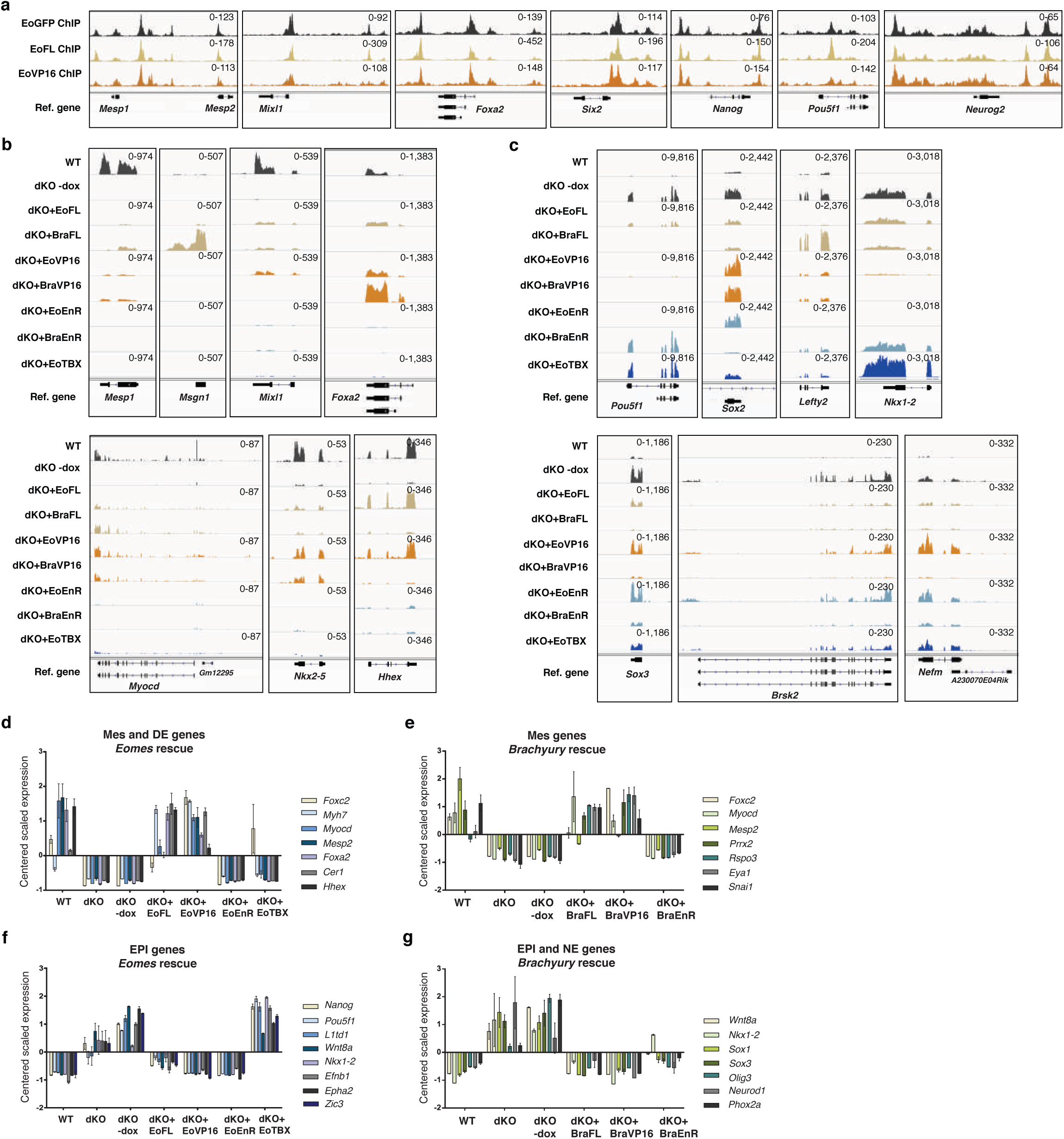
*Eomes*- and *Brachyury*-rescue constructs restore ME program activation and repression of pluripotency and NE programs. **a**, ChIP-seq coverage tracks of EoGFP, EoFL and EoVP16 for indicated genes showing identical binding of GFP and VP16 fusion constructs as the FL EOMES. Counts normalized to RPKM are indicated in the right corner in a-c. **b**, RNA-seq coverage tracks of mesoderm (*Mesp1*, *Msgn1*, *Mixl1*, *Myocd*, and *Nkx2-5*) and definitive endoderm (*Foxa2* and *Hhex*) markers that are rescued by FL and VP16 activator constructs (EoFL, BraFL, EoVP16, and BraVP16) but not by repressor constructs (EoEnR and BraEnR) or EoTBX construct. **c**, RNA-seq coverage tracks of pluripotency (*Pou5f1*, *Sox2*, *Lefty2, and Nkx1-2*) and NE (*Sox3, Brsk2,* and *Nefm*) markers showing selectivity in the transcriptional repression by *Eomes*- or *Brachyury*-FL, VP16, and EnR constructs. Pluripotency genes are predominantly repressed by *Eomes* and NE markers by *Brachyury* in a direct manner as demonstrated by EnR-mediated repression and indirectly by VP16 activator-mediated mechanisms. Expression of EoTBX construct does not rescue the expression of EPI genes. **d**, **e**, Expression levels indicated by centered scaled counts of mesoderm (Mes) and definitive endoderm (DE) marker genes downregulated in dKO cells are rescued after induced expression of *Eomes-* (d) or *Brachyury-* (e) FL, VP16 activator, but not of EnR repressor constructs. Bars represent centered and scaled mRNA expression levels obtained by triplicate experiments of RNA-seq. Error bars indicate SEM in d-g. **f**, **g**, Expression levels indicated by centered scaled counts of EPI and NE marker genes after induced expression of *Eomes-* (f) or *Brachyury*- (g) rescue constructs showing reduced expression by FL, VP16 activator, and EnR repressor constructs, but not with EoTBX construct.

**Supplementary Fig. 6:**
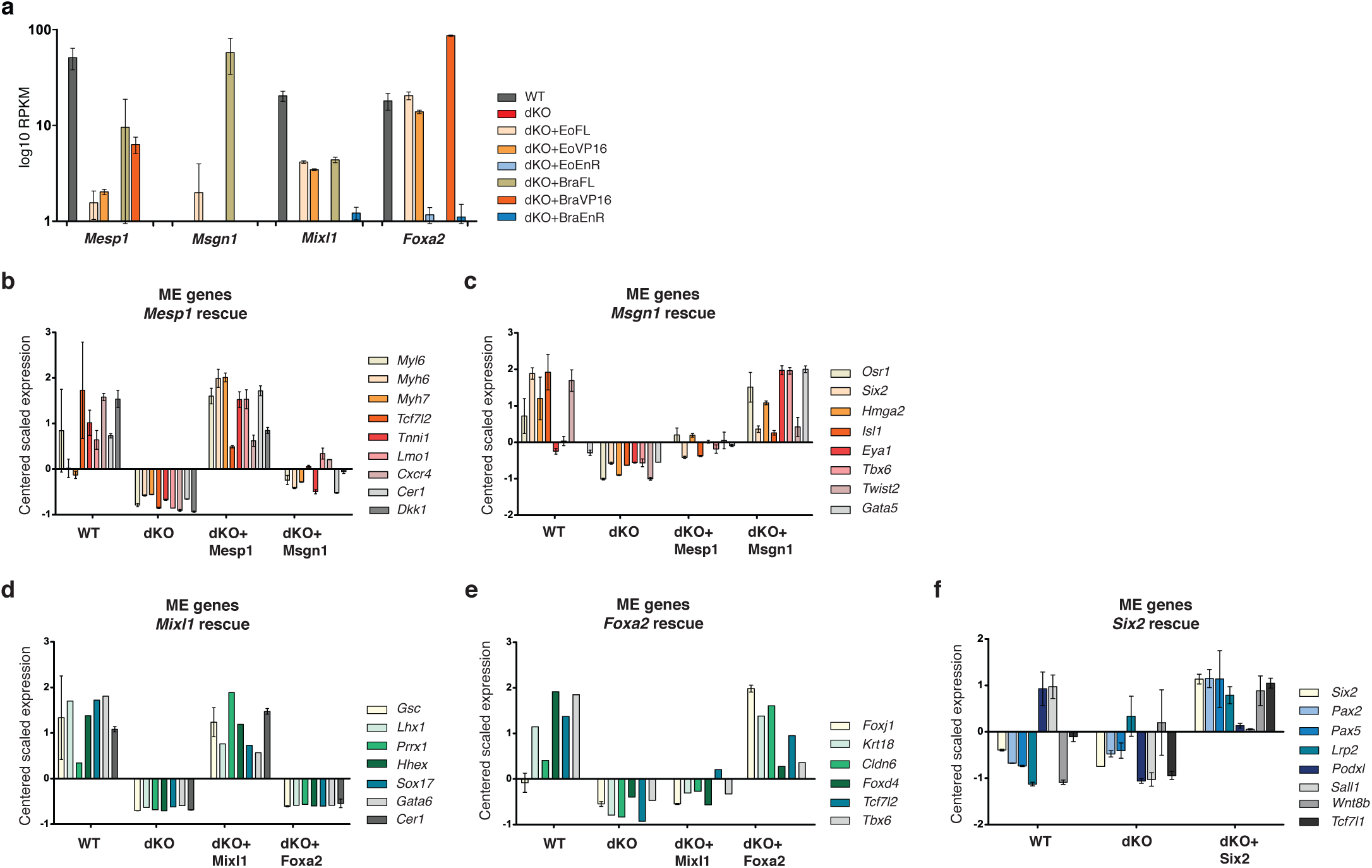
Expression of *Eomes* or *Brachyury* downstream targets *Mesp1*, *Msgn1*, *Mixl1, Foxa2*, and *Six2* in dKO cells activates specific ME gene programs. **a**, Expression levels indicated by log10 RPKM of *Mesp1*, *Msgn1*, *Mixl1*, and *Foxa2* in WT, dKO cells and cells induced with *Eomes-* or *Brachyury-* FL, VP16 and EnR constructs. Error bars represent SEM in a-f. **b-f**, Expression levels of ME genes indicated by centered scaled counts after induced expression of *Mesp1* (b), *Msgn1* (c), *Mixl1* (d), *Foxa2* (e), or *Six2* (f).

## Supplementary Figure: Unprocessed images of Western blots

**Related to Supplementary Figure 1f.**
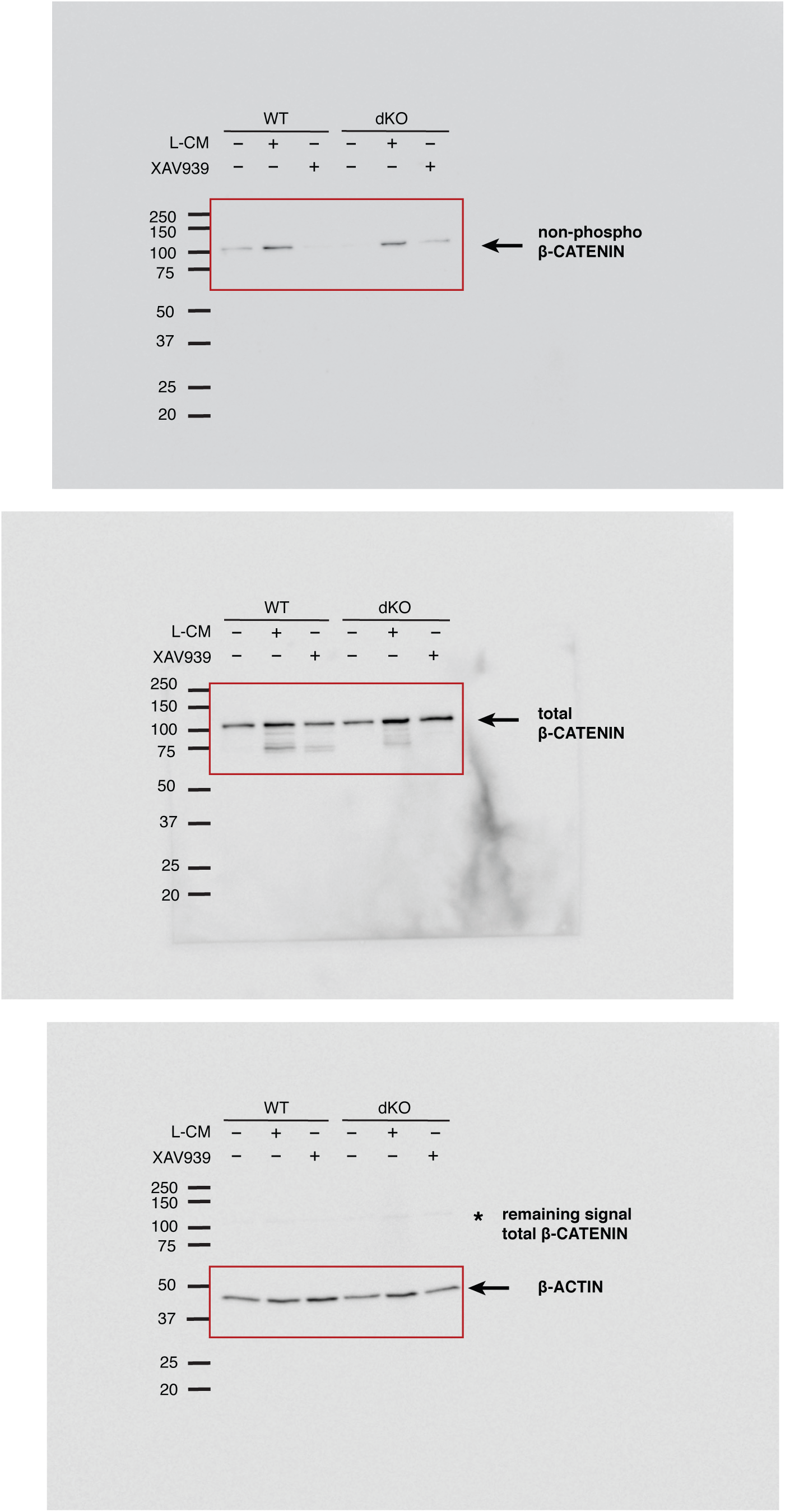

**Related to Supplementary Figure 1g.**
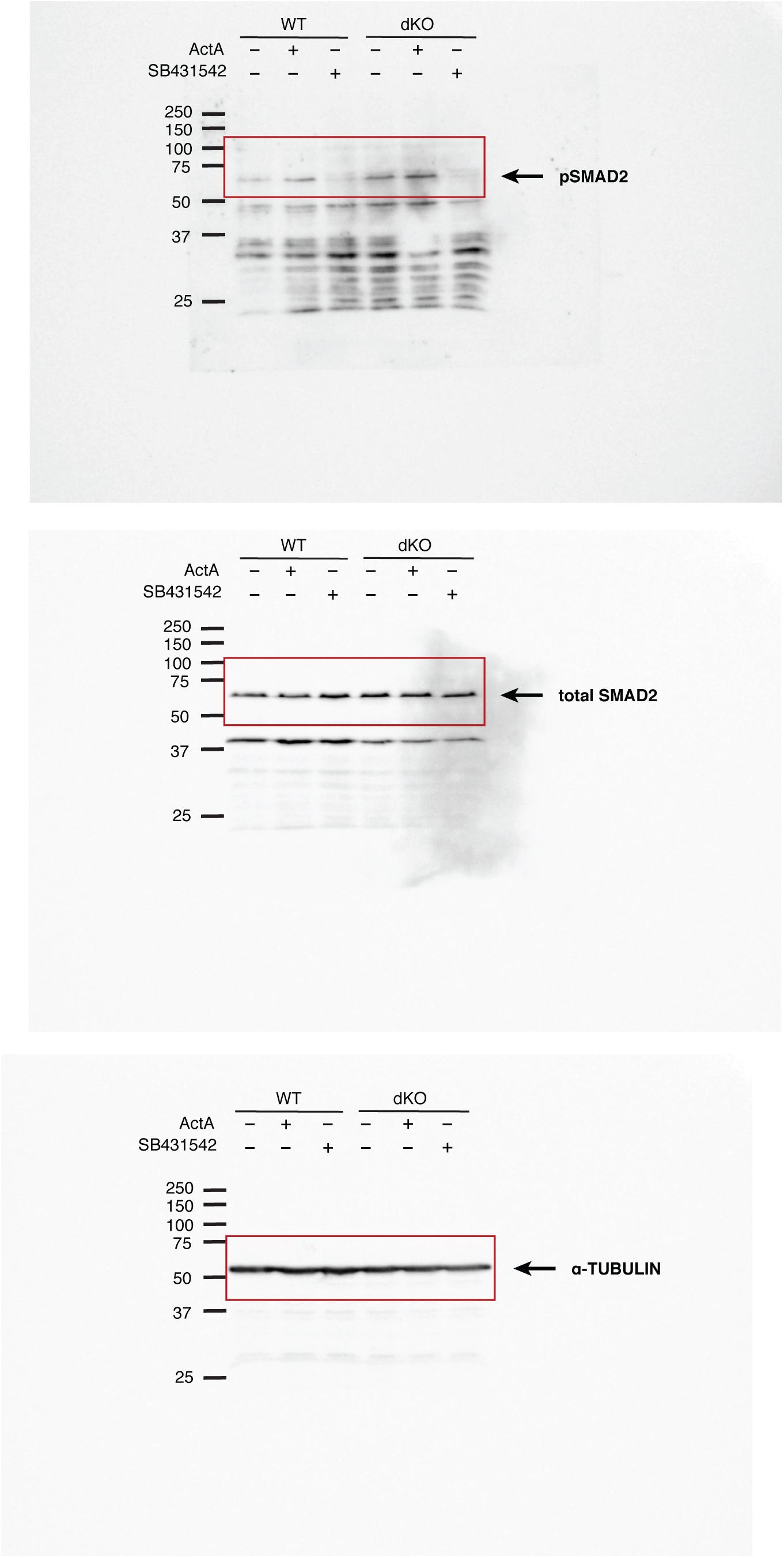

**Related to Supplementary Figure 4b.**
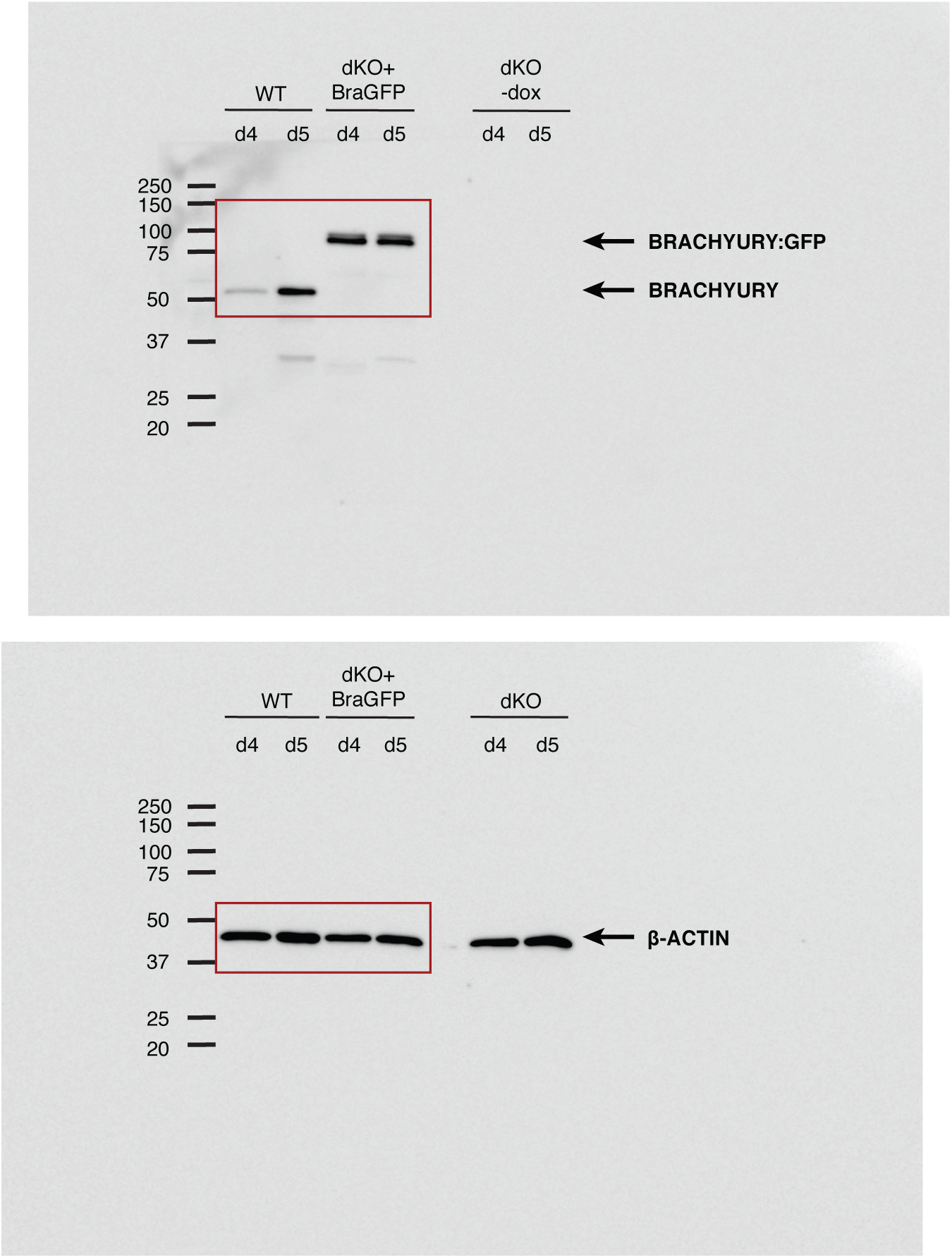

**Related to Supplementary Figure 4c.**
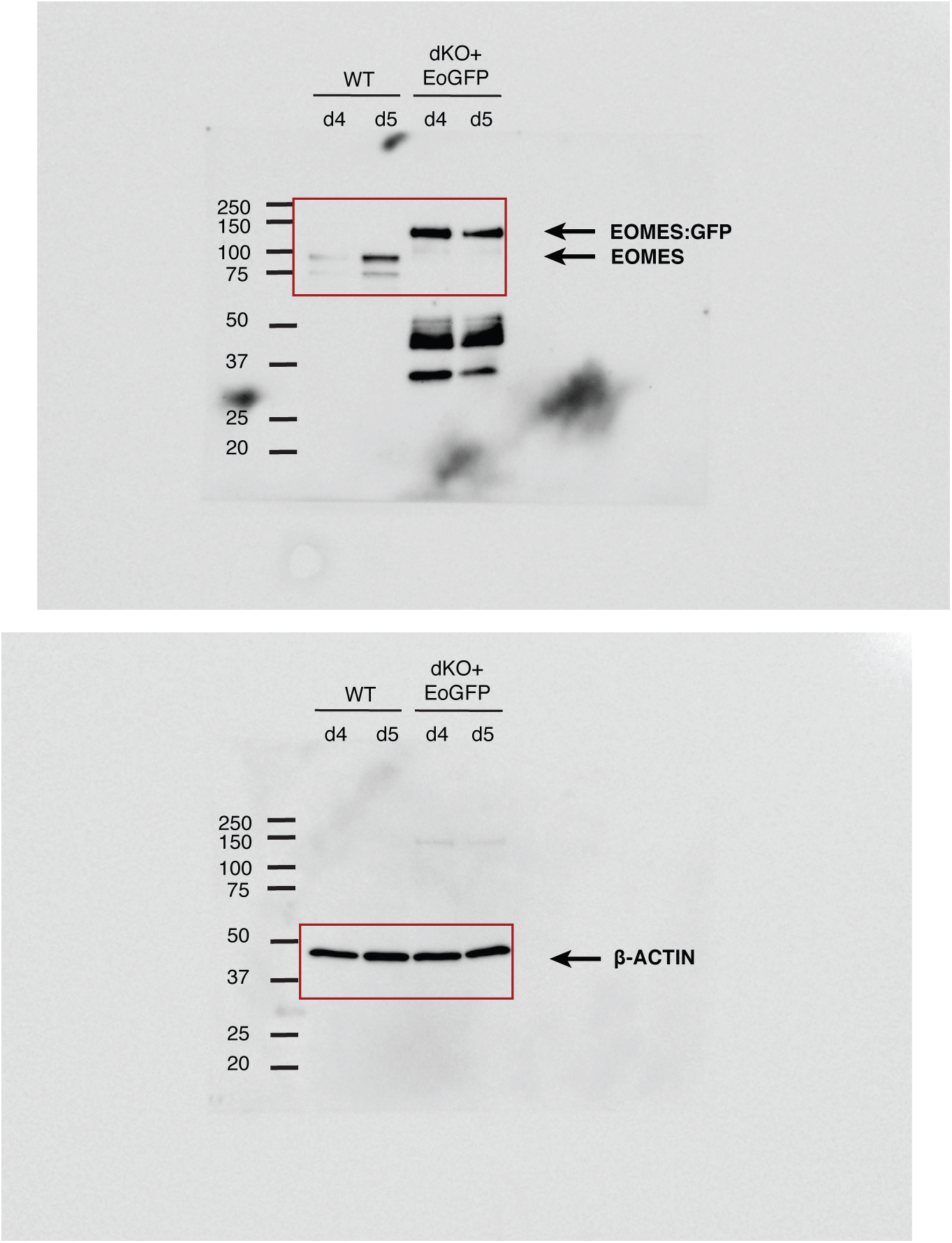

## Methods

### Cell lines

A2lox.Cre mouse embryonic stem cells (ESCs)^1^ were maintained in Dulbecco’s modified Eagle’s medium (DMEM) containing 15% fetal bovine serum (FBS, Gibco), 2 mM L-glutamine, 1X non-essential amino acids (NEAA), 1 mM sodium-pyruvate, 1X penicillin/streptomycin, 100 μM β-mercaptoethanol, Leukemia inhibitory factor (ESGRO LIF, Merck Millipore, 1000 U/ml), and 2i: CHIR99021 (Axon Medchem, 3 μM) and PD0325901 (Axon Medchem, 1 μM) on a monolayer of mitotically inactivated STO mouse fibroblast cells (SNL76/7) or on 0.1% gelatin-coated dishes.

### Animals

*Eomes* and *Brachyury* dKO embryos were obtained by intercrossing *Eomes*^N/+^;Sox2.Cre; *T*^2J/+^ males and *Eomes*^CA/CA^;*T*^2J/+^ females^2–4^. Embryos were dissected at E7.5. For *in situ* hybridization, the *Eomes^Δ^*^Epi^ genotype was inferred by embryonic morphology^2^ (no mesoderm layer, accumulation of cells in the presumptive primitive streak region), while the homozygous *Brachyury*^T2J/T2J^ genotype was identified by the absence of *Brachyury* expression using a *Brachyury* antisense probe. dKO embryos show the characteristic morphology of *Eomes^Δ^*^Epi^ embryos, and lack *Brachyury* expression. For RNA-seq, embryos were genotyped by RT-PCR for expression of *Eomes* and *Brachyury*. Animals were maintained as approved by the local authorities (license number G11/31).

### Generation of Tbx-deficient ESCs

Mouse ESCs deficient for *Eomes* and *Brachyury* were generated using TALEN-mediated gene editing and targeted homologous recombination^5^. In brief, targeting vectors for the integration of fluorescent reporters contain 3’ homology arms of 550-600 bp, including the 3’UTR and the start codon of *Eomes* or *Brachyury* followed by the sequences encoding *Gfp* or *Tomato* reporters, SV40 polyA signal, and loxP-flanked selection cassettes for Neomycin or Hygromycin B. The 5’ homology regions contain 450-500 bp sequences of *Eomes* and *Brachyury* exon 1. A2lox.Cre ESCs were nucleofected simultaneously with TALEN constructs directed to sequences within the first exon of either gene and the respective targeting vector, and resistant colonies screened by PCR for the integration of the fluorescent reporters and frame-shift deletions. ESCs were first targeted at the *Eomes* locus, followed by targeting of the *T/Brachyury* locus. Nucleofection was performed with the Nucleofector Kit for mouse ESC (Lonza), using 2.5 μg of DNA and the Amaxa A-013 program. EoKO cells were selected for 10 days by 350 µg/ml G418 (Invitrogen) or 175 µg/ml Hygromycin B (Thermo Fischer) and single colonies picked, screened by PCR (DreamTaq Green PCR Master Mix, Thermo Fischer, K1082) and expanded. The Cre-mediated deletion of the selection cassettes was induced by treating the cells with 1 μg/ml doxycycline.

### Generation of doxycycline-inducible ESCs

A2lox.Cre mouse ESCs were used to generate dox-inducible gene expression by induced cassette exchange^1^. Prior to nucleofection with the desired p2lox-vector, cells were treated with 1 μg/ml doxycycline (dox; Sigma, D9891-1g) for 1 day to induce *Cre* expression. The cells were nucleofected as described previously and selected with 350 µg/ml G418 (Invitrogen). Correctly targeted clones were screened by PCR, using generic Loxin primers (forward: ATACTTTCTCGGCAGGAGCA, reverse: CTAGATCTCGAAGGATCTGGA) or gene-specific primers in the TRE promoter (forward: ACCTCCATAGAAGACACCG)^6^, and reverse primer in the 5’ end of the inserted gene. For the dox-inducible expression of the inserted gene, 5 μg/ml dox was applied for 2 days in differentiation medium. EoFL, BraFL and EoGFP, BraGFP constructs contained full-length coding regions of *Eomes* or *Brachyury* tagged with the V5-tag sequence or fused via a linker to the *Gfp* coding sequence. The EoTBX DNA binding domain of *Eomes* contained the sequences encoding the amino acids 278-459 of EOMES linked to a V5-tag. Fusions with VP16 and EnR contain the sequences encoding the first 522 amino acids at the N-terminus of EOMES and first 237 amino acids of BRACHYURY, separated by a short linker region. Constructs for the dox-inducible expression of *Mesp1*, *Msgn1*, *Mixl1, Foxa2,* and *Six2* were designed using the full-length coding sequences of corresponding genes.

### ESC differentiation to mesoderm and definitive endoderm

Prior to differentiation, ESCs were depleted of feeders by splitting for 2-3 passages onto 0.1% gelatin-coated 60 mm dishes. For embryoid body (EB) formation, 200 cells in 40 μl ESGRO Complete Basal medium were grown in ultra-low attachment 96-well plates (Greiner BioOne) for 2 days until single EB had formed in every well. EBs were subsequently transferred into 60 mm non-adhesive dishes and allowed to further differentiate in ESGRO Complete Basal medium with 30 ng/ml human recombinant ActivinA (ActA, R&D systems). After 3 days of ActA treatment EBs were plated on fibronectin (20 μg/ml) coated 8-well µ-Slides (Ibidi, 80826) and grown in ESGRO Complete Basal medium supplemented with 5% FBS for additional 2 days.

### ESC differentiation to cardiomyocytes

EBs were formed as described above in 1:1 mixture of DMEM/F-12 medium supplemented with N2 (Gibco) and Neurobasal medium supplemented with B27 (Gibco), with 0.2 mM L-glutamine and 100 μM β-mercaptoethanol in ultra-low attachment 96-well plates (Greiner BioOne). EBs were transferred into 60 mm non-adhesive dishes and treated with 10 ng/ml ActA (R&D systems) from day 2 to day 4, and with 5 μg/ml of dox. Next, EBs were plated on fibronectin (20 μg/ml) coated 8-well µ-Slides (Ibidi, 80826) and grown in IMDM (Gibco) supplemented with 20% FBS, 1% NEAA and 100 μM β-mercaptoethanol for additional 4 days.

### ESC differentiation to neuroectoderm

EBs were formed as described above in ESC medium without LIF and 2i. After 2 days EBs were treated with 100 ng/ml Noggin (R&D Systems) and 10 mM SB431542 (Tocris) for 1 more day with agitation. The next day medium was replaced for ESGRO Complete Basal medium containing 100 ng/ml Noggin and 10 ng/ml ActA^7, 8^.

### qRT-PCR

Total RNA was isolated from approximately 100 EBs (differentiation day 2 and 3) or 25 EBs (differentiation day 4 and 5) using RNeasy Mini kit (Qiagen). 0.5 μg RNA was transcribed to cDNA using QuantiTect Reverse Transcription kit (Qiagen). qRT-PCR was performed using SsoAdvanced Universal SYBR Green Supermix (BioRad) and gene-specific primers (Supplementary Table 9) on a CFX384 Touch Real-Time PCR Detection System (BioRad). TBP (TATA-binding protein) served as a reference gene. The expression was normalized to a reference gene and depicted as a fold change relative to the expression in the undifferentiated ESC. Experiments were performed as biological and technical triplicates, Student’s t-test was used to determine significant differences of the mean values between WT and dKO samples. Asterisks represent p-value - n.s:p>0.05; *:p≤0.05; **:p≤0.01; ***: p≤0.001; ****: p≤0.0001. Error bars indicate SEM.

### Immunofluorescence staining

EBs were fixed for 25 min at room temperature (RT) in 4% PFA, while embryos were fixed overnight at 4°C. The samples were cryo-embedded in 15% Sucrose and 7.5% cold water fish gelatin. Embryos were cut into 8 μm transversal sections and EBs into 10 μm sections. EBs differentiated for 7 days were stained directly in 8-well Ibidi µ-Slides. After permeabilization with 0.2% TritonX-100 in PBS for 20 min at RT and blocking for 2 h with 1% Bovine serum albumin (BSA) in PBS, cells were incubated with primary antibody overnight at 4°C and subsequently with a secondary fluorescence-conjugated antibody and 1:1000 DAPI dilution for 90 min at RT in the dark. The primary antibodies used include: anti-TBR2/EOMES (Abcam, ab23345), anti-BRACHYURY (Santa Cruz, sc-17743), anti-SOX17 (R&D systems, AF1924), anti-FOXA2/HNF3β (Cell signaling, 8186S), anti-FOXC2 (R&D systems, AF6989), anti-NANOG (Thermo Fischer, 14-5761-80), anti-SOX2 (R&D systems, AF2018), anti-OCT3/4 (Santa Cruz, sc-5279), anti-SOX1 (R&D systems, AF3369), anti-TUBULIN β3 (BioLegend, 802001), anti-α-ACTININ (Sigma Aldrich, A7811), and anti-FN1 (Abcam, ab2413). Secondary antibodies used: anti-Rabbit, anti-Goat, anti-Mouse, anti-Rat AlexaFluor 647 (Thermo Fisher), anti-Sheep AlexaFluor 647 (Abcam) and anti-Goat AlexaFluor 488 (Thermo Fisher). Experiments were repeated at least three times and comparable images were taken using same excitation intensity, exposure time and gain values. Confocal imaging was performed using the LSM-I-DUO LIVE 510 META Axiovert microscope equipped with a 40x/1.2 C-Apochromat objective W Korr UV-VIS-IR (Carl Zeiss). Excitation of the fluorophores (DAPI, GFP and Alexa 488, tdTom, Alexa 647) was performed with a two-photon laser at 740 nm, and a single photon laser at 488 nm, 561 nm, and 633 nm, respectively. Plated EBs were imaged using an Axiovert 200M microscope (Carl Zeiss), driven by Visiview (Visitron) imaging software with plan-apochromat objectives, a Yokogawa CSU-X1 spinning disk confocal head with emission filter wheel, and a Coolsnap HQ II digital camera with 488 nm and 640 nm laser lines. Images were processed with Metamorph (Molecular Devices), ZEN (Carl Zeiss) and FIJI (ImageJ)^9^ software.

### *In situ* hybridization

For *in situ* hybridization, embryos were fixed in the deciduae in 4% PFA/PBS overnight at 4°C and embedded in paraffin. Embryos were cut in 8 μm thick transversal sections, deparaffinized, treated for 5min with 1 μg/ml Proteinase K/PBS, fixed for 15 min and incubated in freshly prepared 0.25% acetic anhydride in TEA-buffer for 10 min. The sections were then incubated in hybridization buffer (50% Formamide, 5x SSC, 1% SDS, 50 μg/ml yeast RNA, and 50 μg/ml Heparin in DEPC-H_2_0) containing 10-20 ng/ml digoxigenin-labeled riboprobe, overnight at 68°C. Washes were done the next day with Wash buffer 1 (50% Formamide, 5x SSC and 1% SDS in DEPC-H_2_O) and Wash buffer 2 (50% Formamide and 2x SSC in DEPC-H_2_O) at 65°C and 10 min each wash. Slides were blocked for 30 min in ISH-blocking solution (2% Roche blocking reagent, 1096176, 5% Sheep serum, 1% Tween20 in TBST) and subsequently incubated in antibody solution (5% Sheep serum, 1% Tween20 and 1:2500 dilution of Alkaline Phosphatase-linked α-Digoxigenin antibody, Roche, 11093274910 in TBST) for 2 h at RT. This step was followed by washes with TBST and NTMT buffer (100 mM NaCl, 100 mM Tris-HCl pH9.5, 50 mM MgCa_2_, and 1% Tween20 in H_2_O). BM-Purple (Roche, 1442074) was added to the slides for color development until staining was visible. The samples were fixed with 4% PFA in PBS for 30 min at RT, counter-stained with Eosin for 1 min and mounted in Roti-mount (Carl Roth, HP68.1).

### Dual Luciferase assay

Prior to the experiment 5x10^4^ A2lox or dKO cells were plated on gelatin-coated dishes in 24-well plates in ESC maintaining medium. The next day, medium was changed to Basal medium and cells transfected with Lipofectamine 3000 (Thermo Fischer) according to the manufacturer’s instructions. Transfection was performed with 2.5 μg of pRL-TK plasmid and either pGL4.24-6xARE-Lux^10^, Super 8x TOPflash or FOPflash plasmids (Addgene). After 24h cells were either left untreated in Basal medium or treated with 50 ng/ml ActA or WNT3A/L-cell conditioned medium (L-CM)^11^, and with the inhibitors 10 mM SB431542 (Tocris) or 2 µM XAV939 (Torcis) for the next 24h. Cells were lysed in Passive lysis buffer and the measurement was performed on a TECAN infinite M200 reader with the iControl v1.6 software according to the manufacturer’s instructions. Three biological replicates were quantified by dividing the values measured for the Firefly Luciferase with the Renilla luciferase and normalized to the untreated cells. Student’s t-test was used to determine significance levels between untreated and treated samples. Asterisks represent p-value - n.s:p>0.05; *:p≤0.05; **:p≤0.01; ***: p≤0.001; ****: p≤0.0001. Error bars indicate SEM.

### Western blot

2.5x10^5^ A2lox, dKO, dKO+EoGFP or dKO+BraGFP cells were grown as EBs in suspension culture in 100 mm dishes for 2 days and treated with 30 ng/ml ActA and 5 µg/ml dox from day 2 to day 5 of differentiation. On day 4 and 5 of differentiation, EBs were trypsinized, washed in PBS and lysed in Protein lysis buffer (20 mM Tris-HCl, 137 mM NaCl, 1% NP-40 S Tergitol solution, 2 mM EDTA) containing Complete Protease inhibitor Cocktail (Roche) and Phosphatase inhibitor Cocktail 3 (Sigma Aldrich) for 1h at 4°C on a rotating wheel. For detection of signaling active SMAD2, or *β*-CATENIN cells were grown in Basal medium in 60 mm dishes and treated with either 50 ng/ml ActA (R&D systems) or with WNT3A/L-cell conditioned medium (L-CM)^11^ for 6h, or with the inhibitors 10 mM SB431542 (Tocris) and 2 µM XAV939 (Torcis) for 24h. Lysates were centrifuged at 14,000 rpm, 4°C, 30 min, and supernatant was used to measure protein concentration by the BCA method (Thermo Fischer). 10 or 30 μg of protein lysate was used for each experiment mixed with 1x Sample Buffer (1% SDS, 40 mM Tris-HCl pH 6.7, 5% glycerol, 0.1 % bromophenol blue) freshly supplemented with 100 mM DTT and boiled for 5 min at 95°C. 10% SDS-Polyacrylamide gel was run in 1x Running Buffer (22.54 mM Tris, 172.6 mM Glycine, 0.09% SDS) in Electrophoretic Transfer Cell (BioRad) at 15 mA for approximately 2 h. Proteins were transferred to PVDF membrane in 1x Transfer buffer (47.96 mM Tris, 39 mM Glycine, 0.038% SDS, 20% Methanol) using the Trans-Blot Turbo (BioRad) transfer system with up to 1.0 A; 25 V, 30 min. After transfer, the membrane was blocked in Blocking solution (5% skim milk powder in PBST) for 1 h. Alternatively, for anti-phospho-SMAD2 detection the membrane was blocked in 5% BSA in TBST and washed with TBST. The membrane was incubated with the primary antibody in Blocking solution at 4°C over night. On the following day, membranes were incubated with the secondary antibody in Blocking solution at room temperature for 45 min. For chemiluminescence detection, the membrane was incubated with ECL Prime Western Blotting Detection reagent (GE Healthcare Life Science) and bands were detected using Fujifilm LAS 3000 Image reader. For reprobing of the membrane with different primary antibodies 0,2% NaN_3_ in Blocking solution was used to inactivate Horseradish Peroxidase (HRP). The primary antibodies used include: anti-TBR2/EOMES (Abcam, ab23345), anti-BRACHYURY (R&D Systems, AF2085), anti-phospho SMAD2 (Cell Signaling, 3108S), anti-SMAD2 (BD-Transduction, 610843), anti-non-phospho β-CATENIN (Cell Signaling, 8814S), anti-β-CATENIN (BD Transduction, 610154), anti-*α*-TUBULIN (Abcam, ab4074), and anti-β-ACTIN (Sigma Aldrich, A1978). Secondary antibodies used: anti-Mouse HRP (Dako, P0447) and anti-Rabbit HRP (Dako, P0448).

### RNA-seq

Total RNA from approximately 25 EBs at day 5 of differentiation was isolated using RNeasy Mini kit (Qiagen) and quantified by NanoDrop (Thermo Fisher). Library preparation was carried out using 0.5 μg of RNA with NEBNext Ultra RNA library Prep Kit for Illumina (New England BioLabs, E7530L). For RNA-seq from E7.5 embryos, the embryos were isolated and the epiblast and overlying endoderm layer were dissected from extraembryonic portions along the embryonic-abembryonic border. RNA from the epiblasts and overlying endoderm was isolated using the Qiagen RNeasy Micro kit (74004). Libraries were prepared from 10 ng of total RNA from single embryonic samples with the Ovation SoLo RNA-seq Systems kit (NuGEN, 0501-32). All samples were sequenced in biological triplicates from three age-matched individual embryos. Sequencing was performed at Genomics Core Facility (GeneCore, EMBL, Heidelberg, Germany).

### ATAC-seq

The protocol was modified from Buenrostro et al.^12^ Approximately 100 EBs at day 5 of differentiation were trypsinized to single-cell suspension, washed with PBS, lysed and mixed with 50 µl of transposition reaction mix containing Tagment DNA Enzyme (Nextera DNA Library Preparation Kit, Illumina) and 0.2% Digitonin. The transposition reaction was carried out at 37°C for 30 min, purified using the Qiagen MinElute Kit and eluted in EB-buffer (Qiagen). DNA was amplified by PCR and purified with the Qiagen MinElute Kit (Qiagen, 28004). The libraries were size-selected by polyacrylamide gel electrophoresis (PAGE) by loading the samples onto a 6% polyacrylamide gel and run in TBE buffer (89 mM Tris-HCl, 89 mM Boric acid, 2 mM Na_2_EDTA in H_2_O, pH 8.0). DNA was visualized with SYBRTM^TM^ Gold Nucleic Acid Gel Stain (Invitrogen, 1:10,000 dilution in TBE-buffer). Gel containing fragments between 100-1,000 bp was cut and resuspended in Diffusion buffer (0.5 M CH_3_COONH_4_, 10 mM Mg(CH_3_COO)_2_, 1 mM Na_2_EDTA, 0.1% SDS in H_2_O) in a 1:1 ratio of buffer to gel (mg). After incubation at 37°C for 22 h samples were filtered using a 30 µm CellTrics strainer and purified using the QIAquick Gel Extraction Kit (Qiagen, 28115). DNA concentration was measured with a Qubit Fluorometer (Thermo Fischer, Q32854) and fragment size determined with a Bioanalyzer (Agilent). Libraries were prepared using Nextera DNA Library Preparation Kit (New England BioLabs, E7530L) and sequenced at Genomics Core Facility (GeneCore, EMBL, Heidelberg, Germany). The experiment was performed using biological duplicates.

### ChIP-seq

The protocol was modified from Schmidt et al.^13^, according to Singh et al.^14^ Briefly, ProteinG beads (50 μl per sample) were incubated with 5 μg of Anti-GFP Antibody (Abcam, ab290) or Anti-V5 antibody (BioRad, MCA1360) overnight at 4°C. Starting number of 2.5x10^5^ dKO+EoGFP, dKO+BraGFP, dKO+EoV5 or dKO+EoVP16 cells were grown as EBs in suspension culture in 100 mm dishes for 2 days and treated with 30 ng/ml ActA and 5 µg/ml dox from day 2 to day 5 of differentiation. Uninduced cells were used as negative controls. On day 5 of differentiation, EBs were trypsinized and approximately 3x10^7^ cells was used per ChIP. Cross-linking was performed with 2 mM disuccinimidyl glutarate (DSG, Thermo Fischer, 20593) in the SolutionA (50 mM Hepes, 100 mM NaCl, 1 mM EDTA and 0.5 mM EGTA) for 15 min followed by 1% formaldehyde (Sigma Aldrich, F8775) for additional 10 min. The reaction was stopped by adding 2.5 M Glycine for 5 min. Cells were lysed in Lysis buffer 1 (50 mM Hepes, 140 mM NaCl, 1 mM EDTA, 10% glycerol, 0.5% NP40 and 0.25% TritonX in H_2_O, pH7.5) with protease inhibitors (PI, Complete™ Protease Inhibitor Cocktail, Roche) for 10 min on a rotating wheel at 4°C. Next, pellets were incubated for 5 min at 4°C with Lysis buffer 2 with PI (10 mM Tris, 200 mM NaCl, 1 mM EDTA, 0.5 mM EGTA in H_2_O, pH8), followed by resuspension in Lysis buffer 3 with PI (10 mM Tris, 100 mM NaCl, 1 mM EDTA, 0.5 mM EGTA, 0.1% Na-deoxycholate, 0.5% Lauroylsarcosine in H_2_O, pH8). The DNA was sonicated with the Bioruptor (Diagenode), at high amplitude for 10-12 cycles of 30 s ON/30 s OFF and the fragmentation was evaluated on a 2 % Agarose gel to obtain fragments between 200-650 bp. After sonication, TritonX was added to final concentration of 1%. 10% of the resulting lysate was kept as input control. Beads coupled to antibodies were washed, added to the chromatin and rotated at 4°C overnight. The next day, beads were washed on ice ten times with RIPA buffer (50 mM HEPES, 500 mM LiCl, 1 mM EDTA, 1% NP40 substitute, 0.7% Na-Deoxycholate in H_2_O, pH7.6) and once with TBS. DNA was removed from the beads with Elution buffer (50 mM Tris, 10mM EDTA, 1% SDS in H_2_O, pH8) for 6-18 h at 65°C. The supernatant was mixed with one volume of TE buffer (10 mM Tris, 1 mM EDTA in H_2_O, pH7.4), incubated with RNAse A at 37°C for 1 h, and ProteinaseK at 55°C for 2 h. Phenol-chloroform extraction of the DNA was performed using GlycoBlue co-precipitant (Invitrogen, AM9516) to visualize the pellet and 5PRIME Phase Lock tubes to facilitate the separation of the phases. The pellet was resuspended in 50 μl of 10 mM Tris-HCl pH8. Libraries were prepared with NEBNext Ultra II DNA Library Prep Kit for Illumina (NEB, E7645S), size selected on a 1.3% agarose gel by cutting out fragments between 200-650 bp of size and extracted from the gel using QIAquick gel extraction kit (Qiagen, 28704). Samples were sequenced at Genomics Core Facility (GeneCore, EMBL, Heidelberg, Germany). Biological duplicates were used for ChIP-seq experiments.

### H3K27ac ChIP-seq

Around 100 WT and dKO EBs after 5 days of differentiation were used for H3K27ac ChIP-seq experiment. The procedure was performed with ChIP-IT High Sensitivity kit (Active motif, 53040). 5 μg of Anti-H3K27ac antibody (Abcam, ab4729) was used. Libraries were prepared with NEBNext Ultra II DNA Library Prep Kit for Illumina (NEB, E7645S) and size selection was performed as described above. Samples were sequenced at Genomics Core Facility (GeneCore, EMBL, Heidelberg, Germany).

## Statistics and reproducibility

### RNA-seq

Sequenced reads were mapped to the mouse reference genome GRCm38/mm10 (iGenomes, Illumina) containing the chromosomes 1-19, X, Y and M using Rsubread v1.28.1 package in R v3.4.3^15^. The genomic annotations used for the mapping, counting and downstream analyses were contained in gtf files provided by the iGenomes reference genome bundles (archive-2015-07-17-32-40 and archive-2015-07-17-33-26 for GRCM38 and mm10, respectively). Genomic features were counted using the Rsubread::featureCounts function^15^. Differential expression analysis was carried out using DESeq2 v1.18.1 in R^16^, which applies Negative Binomial GLM fitting and Wald statistics on the count data. The results were filtered for adjusted p-value<0.05 and log_2_FC as indicated in the figure legends. Prior to fitting and statistical analysis, counts were normalized by library size (DESeq2::estimateSizeFactors) and gene-wise dispersion (DESeq2::estimateDispersions) to normalize the gene counts by gene-wise geometric mean over samples. For plotting the heatmaps, gene-wise scaling was performed on the normalized counts to cluster the counts for the expression tendency between the samples (stats::kmeans function). In addition, the counts were centered by setting the gene-wise mean to zero, which explains why some count values are negative (below gene-wise mean). Visualization of the clustered data was performed using the pheatmap::pheatmap function (pheatmap v1.0.10) with deactivated function-intrinsic clustering. For the visualization of coverage tracks in the Integrated Genome Viewer (IGV) v2.3.93^17^, mapping was performed on Galaxy platform^18^ using Bowtie2 (v2.3.4)^19^ on mm10 reference genome with default settings. Obtained bam files from biological triplicates were merged using Merge BAM files tool in Galaxy (picard v1.56.0) and the duplicates were removed by RmDup tool (samtools v1.3.1)^20^. Coverage files were created using bamCoverage tool (deepTools v3.0.2)^21^ with bin size 10 bases and reads per kilobase per million (RPKM) normalization. The Group Autoscale function was used to scale the tracks in IGV^17^. Jitter plots were produced by plotting the centered scaled counts for selected gene clusters by using ggplot2::geom jitter function (v3.0.0) in R-v3.4.4. Centered and scaled counts for selected genes were represented as bar charts for rescue experiments by dox-inducible expression of various constructs. These bar charts were created with GraphPad Prism software v5.04, where error bars indicate SEM between 3 biological replicates.

### Gene Ontology overrepresentation

For GO-term analysis, gene names were converted to Entrez IDs (biomaRt v2.34.2,org.Mm.eg.db v3.5.0, and clusterProfiler v3.6.0; clusterProfiler::bitr function converting ALIAS to ENTREZID with org.Mm.eg.db as database), and the enrichment of gene groups assessed using the clusterProfiler::enrichGO function (GO-terms from org.Mm.eg.db)^22^. Biological pathway enrichments were assessed using the clusterProfiler::enrichKEGG function with pathway annotations from KEGG.db v3.2.3. Redundant GO-terms were removed using the clusterProfiler::simplify function (cut-off 0.7 and by=p.adjust). The parameters for these enrichment functions: p-value and q-value cut-off of 0.05 and the pAdjustMethod Benjamini-Hochberg.

### Principal component analysis

Principal component analysis was performed using ClustVis online tool^23^ with normalized reads of 3 biological replicates using differentially expressed genes obtained by DEseq2. Unit variance scaling was applied to rows. Singular value decomposition with imputation was used to calculate principal components.

### ATAC-seq

Reads were mapped to the mm10 genome using Galaxy platform Bowtie2 v2.3.4^19^ with default settings. Results from biological duplicates were merged for subsequent analysis using Merge BAM files (picard v1.56.0). After removing duplicates with RmDup (samtools v1.3.1)^20^, peaks were detected using MACS2 (v2.1.1.20160309.4)^24^ with BAMPE format of the input file, default settings for building the shifting model (confidence enrichment ratio against background 5-50; band width 300) and minimum q-value cut-off for detection of 0.05. For visualization in IGV, the coverage files were created using bamCoverage (deepTools v3.0.2)^21^ with bin size 10 bases and normalization to RPKM.

### ChIP-seq

Processing of reads until MACS2 peak detection^24^ was performed as described in the ATAC-seq section. Input served as control for MACS2 peak calling. Peaks were further filtered for the pile-up value (PU, height of the peak at the summit, provided in the Peaks tabular file from MACS2) PU>120 for EoGFP, PU>40 for BraGFP, PU>60 for EoV5, and PU*≥*36 for EoVP16 ChIP-seq. Peaks detected by MACS2 in the input control and dKO-dox control were subtracted from the genomic intervals detected in ChIP-seq treatment file using SubtractBed tool (bedtools v2.27.0)^25^ in Galaxy, where entire features were filtered out if the overlap of minimum 1 bp was found. For visualizations of read coverage in IGV^17^, coverage files were created using the bamCoverage tool (deepTools v3.0.2)^21^ with bin size 10 bases and normalization to RPKM. The coverage track for EO+BRA ChIP was created by Combine Data Track tool with setting Operation:add in IGV. The peak detection by MACS2 is generated as bed file by adding the processed intervals from EoGFP and BraGFP experiments and combining overlapping intervals into a single interval using MergeBED (bedtools v2.27.0)^25^ function on Galaxy. Overlapping intervals from ChIP-seq peak files with ATAC-seq peaks were generated using Intersect intervals of two datasets tool (bedtools 2.27.0)^25^. The genomic intervals found in this intersection (bin size 50) associated with upregulated or downregulated genes in dKO were centered to the middle of the interval considering 2.5 kb upstream and downstream, sorted in the descending order and plotted. Heatmaps were made with ATAC-seq or H3K27ac ChIP-seq normalized counts to RPKM depicted as the intensity of red color, while the peak profile was created with averaged RPKM values using the computeMatrix and plotHeatmap functions of deepTools (v.3.0.2)^21^.

### Genomic Regions Enrichment of Annotations

Statistical enrichment for association between genomic regions detected as ChIP-seq peaks after MACS2 peak calling and filtering, and annotations of putative target genes was performed using Genome Regions Enrichment Annotations Tool online tool (v3.0.0)^26^ using Basal plus extension setting. Each annotated gene was assigned a basal regulatory region of 5 kb upstream and 1 kb downstream of the transcription start site, and the regulatory domain was extended in both directions up to 1,000 kb until the nearest gene’s basal domain.

### Genomic distribution pies

Pie plots were generated by the ChIPseeker::plotAnnoPie function (ChIPseeker v1.14.2)^27^ after annotation by the ChIPseeker::annotatePeak function. Gene and transcript annotations were used from the packages org.Mm.eg.db (v.3.5.0) and TxDb.Mmusculus.UCSC.mm10.knownGene (v3.4.0), respectively.

### Motif enrichment analysis

Homer (v4.9.1)^28^ motif analysis was performed on bed files generated by MACS2 peak-caller on the reference genome mm10 with default settings for the basic findMotifsGenome.pl command. The motif enrichment for WT ATAC-seq peaks was calculated using dKO ATAC-seq peaks as a background and *vice versa*.

### Venn diagrams

Proportional Venn diagrams to show unique and intersecting elements of gene lists were generated using the web-tool available at http://bioinformatics.psb.ugent.be/webtools/Venn/.

### Data availability

The accession number for the primary sequencing data acquired in this paper is GSE128466. RNA-seq data of mouse EpiSC is available at GSE99494, ATAC-seq of mouse ESCs at GSE94250 and ATAC-seq of mouse EpiSCs at GSE110164. All other data are available from the authors on reasonable request.

